# Gene regulatory network analysis predicts cooperating transcription factor regulons required for FLT3-ITD+ AML growth

**DOI:** 10.1101/2023.07.18.549495

**Authors:** Daniel J.L. Coleman, Peter Keane, Rosario Luque-Martin, Paulynn S Chin, Helen Blair, Luke Ames, Sophie G. Kellaway, James Griffin, Elizabeth Holmes, Sandeep Potluri, Salam A. Assi, John Bushweller, Olaf Heidenreich, Peter N. Cockerill, Constanze Bonifer

## Abstract

AML is a heterogenous disease caused by different mutations. We have previously shown that each mutational sub-type develops its specific gene regulatory network (GRN) with transcription factors interacting with multiple gene modules, many of which are transcription factor genes themselves. Here we hypothesized that highly connected nodes within such networks comprise crucial regulators of AML maintenance. We tested this hypothesis using FLT3-ITD mutated AML as a model and conducted an shRNA drop-out screen informed by this analysis. We show that AML-specific GRNs predict identifying crucial regulatory modules required for AML but not normal cellular growth. Furthermore, our work shows that all modules are highly connected and regulate each other. The careful multi-omic analysis of the role of one (RUNX1) module by shRNA and chemical inhibition shows that this transcription factor and its target genes stabilize the GRN of FLT3-ITD AML and that its removal leads to GRN collapse and cell death.

## Introduction

Cancer occurs when mutations in signalling genes, transcription factors (TFs) and epigenetic regulators cause a block in the normal program of differentiation and an increase in proliferation. Acute Myeloid Leukaemia (AML) is no exception and, due to the clonal nature of the disease, different driver mutations of the cancer cause distinct patterns of gene regulation in AML cells^1,2^. Due to the differing effects of these mutations, there is a drive to identify druggable targets for each distinct AML subtype to tailor treatment to the individual and reduce the need for intensive chemotherapy which is often not tolerated by elderly patients.

AML with an internal tandem duplication of the FLT3 receptor (FLT3-ITD+ AML) which converts a ligand responsive receptor into a constitutively active molecule is a highly aggressive AML subtype which is frequently refractory to first line therapy^3^. The FLT3-ITD mutation occurs on the background of other mutations, mostly in epigenetic regulators such as DNMT3 or TET2, or as a result of clonal evolution^4,5^. For reasons that are as yet unknown, this mutation often occurs together with a mutation in nucleophosmin (NPM1). Less than 60% of FLT3-ITD AML patients over 60 year-olds reach complete remission and patients often relapse within a year with an overall relapse rate of 77% ^6^. For this reason, therapy efforts have focussed on the development of improved FLT3-specific inhibitors, resulting in the approval of Gilteritinib for use in a clinical setting. However, relapse still frequently occurs after treatment with these inhibitors, often due to activating mutations in other signalling genes ^7^. The identification of other druggable targets in FLT3-ITD AML is therefore necessary to improve the outcome of patients. To this end, various efforts have been conducted to employ genome-wide RNAi or CRSPR/Cas9 screens which highlighted numerous targets ^8,9^. However, a caveat of such screens is often that they come up with a large number of targets, many of which are also important for normal cells, and often require follow-up experiments to identify therapeutic windows for their inhibition. To home in on the true AML subtype-specific targets, it is therefore necessary to elucidate the fine details of the molecular mechanisms driving AML growth and maintenance of each sub-type.

Differential gene expression and thus cellular identity are controlled by the regulatory interactions between transcription factors (TFs) and their target genes which form extensive interacting gene regulatory networks (GRNs). We have previously shown that AML blast cells with FLT3-ITD and FLT3-ITD/NPM1 mutations display a specific chromatin signature distinct from healthy CD34+ cells^1^. To construct GRNs, we then integrated transcriptomic (RNA-Seq), Hi-C and digital footprinting data based on high resolution DNaseI-seq experiments^2^. The comparison between the GRNs of normal and malignant cells identified TF families showing FLT3-ITD+ AML subtype-specific interactions with their targets which could be attributed to aberrant expression of genes promoting AML survival. For several of these TFs, such as the AP-1 TF family we could indeed show that they were required for the maintenance of AML but not normal cells. For example, inhibiting AP-1 activity completely blocked tumorigenesis in two different patient-derived xenograft (PDX) mouse models^2^. However, the molecular details of how specific TFs contribute to AML development and their interaction with genes encoding other TFs is largely unclear.

To answer these questions, we generated a refined GRN for FLT3-ITD and FLT3-ITD/NPM1 AML. Based on this analysis, we designed a targeted shRNA screen to interrogate the contribution of specific TF network nodes and their down-stream targets to AML establishment and maintenance. These analyses highlight a crucial role of several different TFs including RUNX1 in FLT3-ITD pathology and identify drug responsive AML-subtype-specific and overlapping regulatory modules. Taken together, our data show that identifying AML-subtype specific GRNs is predictive for genes required for AML maintenance.

## Results

### Constructing a refined FLT3-ITD+ AML GRN

Cell type-specific gene expression is largely encoded in the distal enhancer elements physically interacting with their cognate promoters^10^. Our previously constructed gene regulatory networks for AML with FLT3-ITD and FLT3-ITD/NPM1 genotypes highlighted interactions between TF families and distal elements demonstrating that each AML-sub-type displayed a specific gene regulatory phenotype distinct from that of normal cells. However, to demonstrate the full regulatory relevance of the GRN and its downstream targets we developed a novel computational method to produce a GRN that contained the full set of regulatory interactions between all cis regulatory elements including promoters as depicted schematically in Figure 1A (see methods). Briefly, open chromatin regions specific to AML (>3xfold change (FC) as compared to normal cells) were determined using high-read depth DNAseI-seq data highlighting occupied motifs assigned to a transcription factor family^2^ as determined by digital footprinting ^11^. Such regions were assigned to promotors using previously determined promoter-capture HiC (CHiC) interactions from a FLT3-ITD+ AML sample or the nearest gene (both within 200kb of the promoter)^2^. RNA-seq was used to identify genes specifically upregulated in the FLT3-ITD+ AML subtype compared to healthy CD34+ cells (>2 FC, p<0.1). These data were plotted as a GRN showing specific interactions between TF families and their target genes in FLT3-ITD+ AML, which each TF interacting with its own specific gene modules of down-stream target genes. These modules can be AML-specific, shared or specific only to normal cells (Fig 1A, bottom panel).

**Figure 1.**
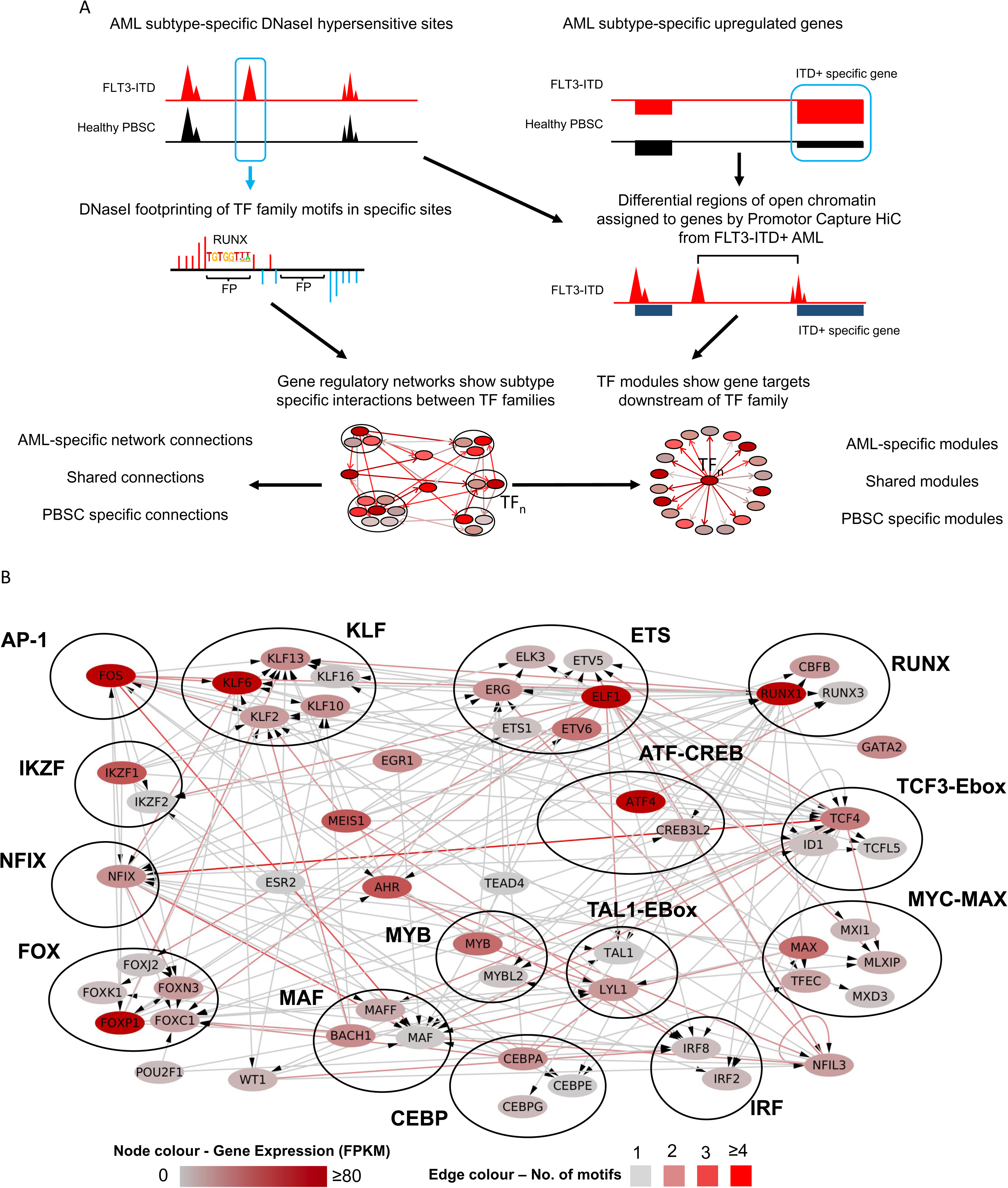
A refined gene regulatory network for FLT3-ITD and FLT3-ITD/NPM1 AML. A: Scheme of transcription factor (TF) network and regulatory module generation. B: FLT3-ITD/FLT3-ITD NPM1 TF network generated by integrating data from DHSs specific for AML identified in the union of DHSs from 9 patients as compared to healthy PBSCs. TF families binding to the same motif are encircled. The colour code for nodes and edges is explained at the bottom of the figure. Edges indicate the presence of a TF motif in a region of chromatin open in FLT3-ITD AML and assigned to the target gene by HiC or nearest gene (<200 kb). Nodes represent individual TF genes within a family. Only genes with connections from other TF families are shown, if there are no incoming connections from other TF families then the highest expressed gene in the TF family in FLT3-ITD AML is shown.

The majority of FLT3-ITD mutations co-occur with the NPM1 mutation ^12,13^ which was also true for our patient cohort (Table 1). We have previously shown that FLT3-ITD and FLT3-ITD/NPM1 cluster together ^2^ and form very similar AML-specific GRNs (Figure S1A). We therefore merged the data from both mutant groups and constructed a shared GRN (Figure 1B) with a large number of highly connected nodes of AML-specifically upregulated TF genes such as the KLF, RUNX, C/EBP, FOX families and NFIX, all displaying multiple edges being connected to other TF encoding genes. This analysis confirmed that RUNX, ETS and AP-1 motifs were highly enriched in FLT3-ITD-specific DHSs and occupy a central position in the network with multiple edges going in and out.

### The refined FLT3-ITD GRN predicts genes required for AML maintenance

We next tested the hypothesis that highly connected nodes within the refined GRN and some of their targets would be important for the maintenance of FLT3-ITD AML. To this end, we performed an informed shRNA screen targeting 161 genes selected from the GRN and the TF modules ^14^ (Fig 2A). The screen was performed in two FLT3-ITD cell lines (MV4-11, MOLM14) *in vitro*, and for MOLM14 in vivo using NSG mice as summarised in Fig 2A.

**Figure 2.**
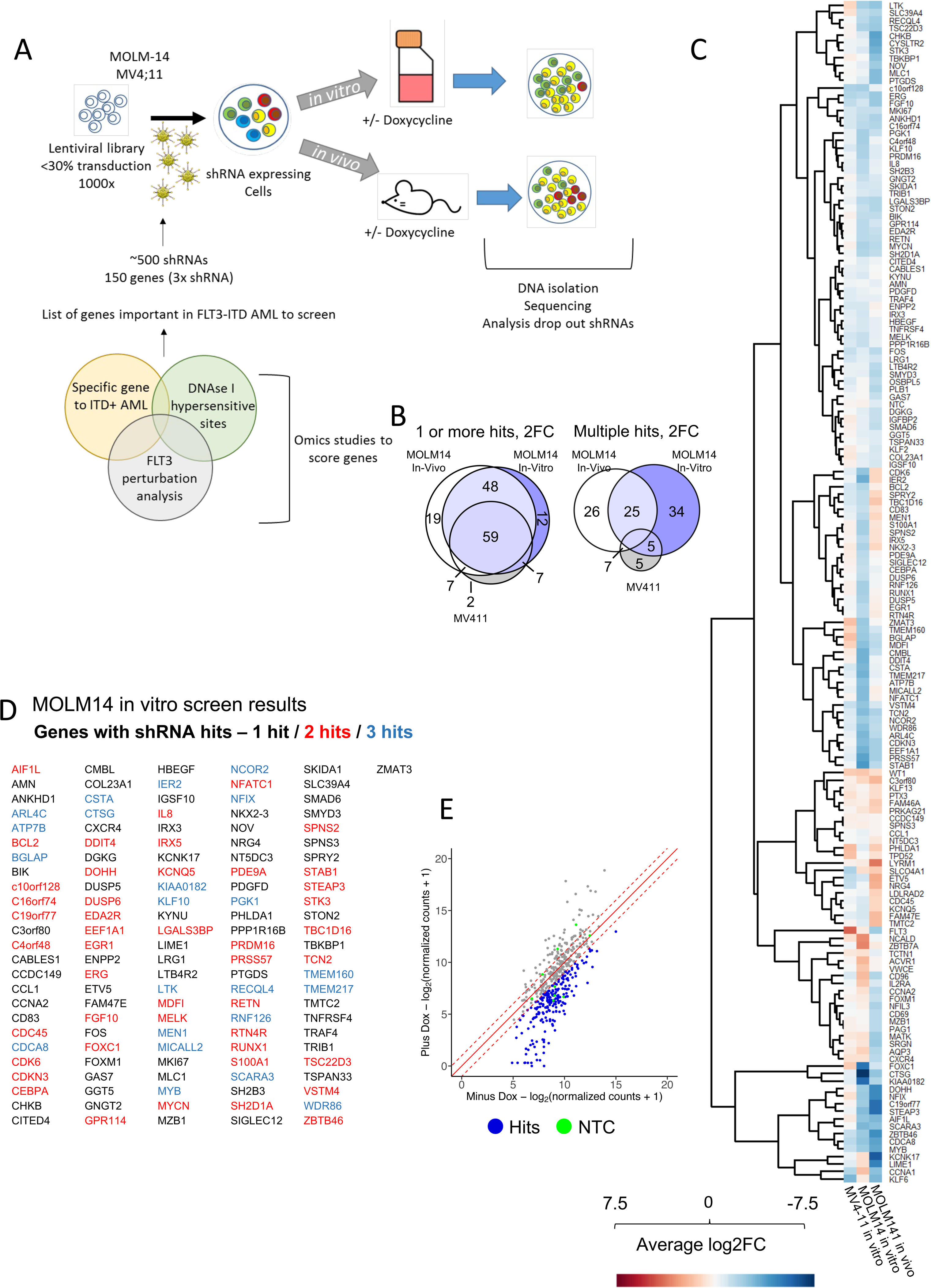
The GRN of FLT3-ITD and FLT3-ITD/NPM1 informs a highly efficient shRNAi screen. A: Scheme of shRNA screen *in vitro* and *in vivo* strategy. B: Venn diagrams of genes with lost shRNAs in screen – 2 FC, 1 or more hits – and genes with multiple hits showing a 2 FC in abundance. C: Average fold-change of shRNA abundance in three screens (average of all 3 shRNAs per gene) with the target genes plotted on the right. D: MOLM14 *in vitro* screen results, list of genes with shRNA hits. E: shRNA abundance after screen in MOLM14 *in vitro* plotted as scatter plot.

To identify a subset of genes specifically important in FLT3-ITD/NPM1 mutated AML survival genes we conducted a rigorous filtering strategy (Fig S1B). Potential targets are listed in Fig S1C and were scored by their specific RNA expression in FLT3-ITD AML vs PBSCs, whether DHSs linked to the gene by CHiC were specific to FLT3-ITD+ AML and contained motifs relating to key nodes in the GRN. We also included genes which were repressed after treatment with FLT3i in a PDX generated from a primary AML *in vitro.* Each target was covered by 3 shRNAs as well as 6 non-targeted control shRNAs and was cloned into a DOX-inducible lentiviral vector expressing a constitutive Venus and an inducible dTomato fluorochrome (Fig S1C). In our previous studies we had already validated the importance of FOXC1, NFIX and the AP-1 family represented by *FOS* ^2^ which were included as positive controls. Our targeted depletion screen showed a significant overlap between targets (Fig 2B), and, importantly, a very high hit-rate, with the majority of selected targets being depleted and nearly 50% of them being important for growth *in vitro* and *in vivo* (Figure 2B-E; Figures S2 A-D). Genes depleted in the screen included multiple TF genes (*EGR1, FOS, RUNX1, CEBPA, KLF2* and *FOXC1*, together with cell cycle regulators, epigenetic regulators and signalling proteins. We selected shRNAs against the TF genes *NFIL3, RUNX1* and *EGR1* to manually validate the results in the cell lines using colony assays in MOLM14 cells (Fig S2 E-G). We also expressed a dominant negative version of C/EBP ^15^ from a doxycycline inducible plasmid, to highlight the importance of this TF family for FLT3-ITD colony forming ability. Our screen also identified the Dual-specificity phosphatases DUSP5 and DUSP6 as hits. To validate a non-TF target, we treated various cell lines representing different AML sub-types with the DUSP inhibitor BCI (Fig S2i) and showed that it is an efficient inhibitor of cell growth, albeit not in a FLT3-ITD-specific fashion.

Taken together, our experiments demonstrate that constructing a disease-specific GRN based on primary AML cell data is highly predictive for identifying network nodes required for the growth of AML cell lines and primary AML but not healthy cells.

### Identification of FLT3-ITD specific overlapping transcription factor modules

Since TFs and other regulators of gene expression are part of an interacting network, targeting such molecules in a clinical setting requires careful examination of the effects of perturbation on their down-stream targets and how GRNs shift in the absence of specific TFs. We therefore connected the above-described transcription factors to the wider patient derived GRN by using digital footprinting and (where available) ChIP analyses to identify AML-specific and shared TF modules. These analyses were conducted for individual TFs (C/EBP, RUNX, EGR) and the previously studied TF families AP-1, FOXC1, NFIX) which were linked to down-stream genes specifically up-regulated in FLT3-ITD/NPM1 cells as outlined in Fig 3A). We also included occupied binding sites shared with normal cells as they often contain motifs for signalling-responsive TFs which could stimulate AML-specific gene expression in an aberrant signalling environment. Fig S3A shows that each TF is connected to a large number of up-regulated target genes, many of which are shared between different modules as exemplified by *DUSP5/DUSP6* or *WT1* (highlighted), suggesting that such genes are regulated by more than one factor. We also observed cross-regulation between modules as exemplified by FOXC1 being part of the NFI module (highlighted). Note that FOXC1 and NFIX are examples of TFs which are aberrantly expressed as compared to healthy cells ^2,16^. FOXC1 is truly mis-expressed, and NFIX plays a role in hematopoiesis and HSC survival ^17,18^ but is strongly up-regulated in FLT3-ITD/NPM1 cells. Their expression is therefore a part of the aberrant FLT3-ITD/NPM1 leukemic phenotype.

**Figure 3.**
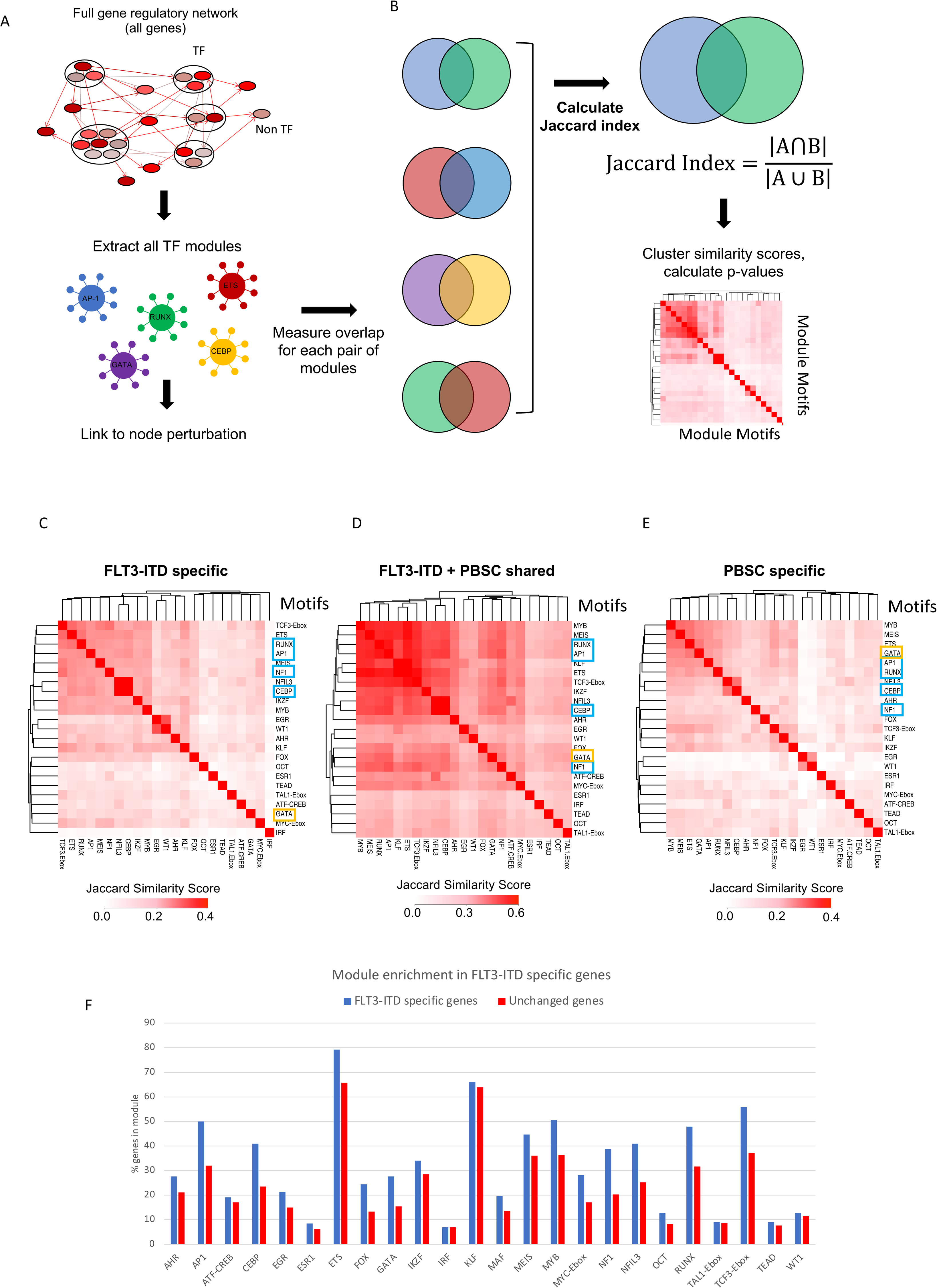
Identification and comparison of regulatory TF modules. A: Scheme of module identification. TFn: Transcription factor. B: Scheme of identifying Jaccard similarity of TF regulatory modules in FLT3-ITD specific DHSs compared to PBSCs. C: Jaccard similarity of TF regulatory modules in FLT3-ITD specific DHSs as compared to PBSCs. D: Jaccard similarity of TF regulatory modules in DHSs shared with PBSCs. Blue boxes: TFs whose module was perturbed in this study. Yellow box: GATA motifs. E: Jaccard similarity of TF regulatory modules in PBSC specific DHSs as compared to FLT3-ITD. F: Enrichment of modules in FLT3-ITD in sites linked to upregulated mRNAs compared to PBSCs. Control are genes that are at the same level of expression in PBSC and FLT3-ITD AML. The y axis shows the percentage of genes in each group which are assigned to a TF family module.

To examine the degree by which nodes are shared between AML and normal cells we calculated their similarities as shown in Fig 3B. This analysis showed that the FLT3-ITD-specific GRN (Fig 3C) contains a central cluster of overlapping nodes for the TF modules TCF3-Ebox, ETS, RUNX, AP-1, MEIS, NFI, C/EBP/NFIL3, IKFZ and MYB which is also found in the shared sites (Fig 3 D). The PBSC-specific central module cluster (Fig 3E) contained similar motifs but is characterised by the additional presence of GATA motifs indicating a more immature state of cells. We next determined all modules of the whole GRN and asked the question which ones were enriched in FLT3-ITD specifically expressed genes (Fig 3 F). This analysis showed again that specific modules were overrepresented by FLT3-ITD-specifically expressed genes, including again AP-1, C/EBP, NFI and RUNX1 modules.

Taken together, this analysis showed that FLT3-ITD specifically expressed genes are regulated by specific and overlapping sets of TF modules.

### Perturbation of FLT3-ITD specific TF modules highlights regulatory relationships based on combinatorial TF action

We next examined the crosstalk between selected TF modules by genome wide analyses in primary AML cells. To this end, we interrogated the RUNX1 and NFIX modules by shRNA and the larger C/EBP and AP-1 families by expressing their dominant-negative counterparts, followed by ATAC-Seq and RNA-Seq (Fig S4, B-F). As controls, we tested these targets together with shRNA against *DUSP5, FOXC1* and *EGR1* in healthy CD34+ PBSCs by transducing a mini-library of lentivirally expressed shRNAs together with a non-targeting control (NTC), where no suppression of growth was observed (Fig S4A). In addition, we transduced AML cells with vectors expressing a DOX-inducible dominant negative peptides targeting the AP-1 and dnC/EBP TF families followed by a colony assay (Fig S4B-C). The inactivation of transcription factors had a profound effect on the growth of AML but not healthy cells. In contrast, targeting DUSP5 and 6 phosphatase activity with inhibitor BCI affected the growth of both AML and healthy cells, albeit with different efficiency as shown by inhibitor experiments (Fig S4D)

The analyses of ATAC-Seq data showed that each perturbation had a strong effect on the chromatin landscape with thousands of sites opening and closing, indicating that factor perturbation not only affected growth, but also changed the identity of cells (Figs 4A-D). However, despite sites being gained or lost, with one exemption which is described below, the enriched motif compositions in those sites did not significantly change, with PU.1, RUNX, C/EBP and AP-1 motifs dominating the picture, suggesting that the system rewires using the same modules.

**Figure 4.**
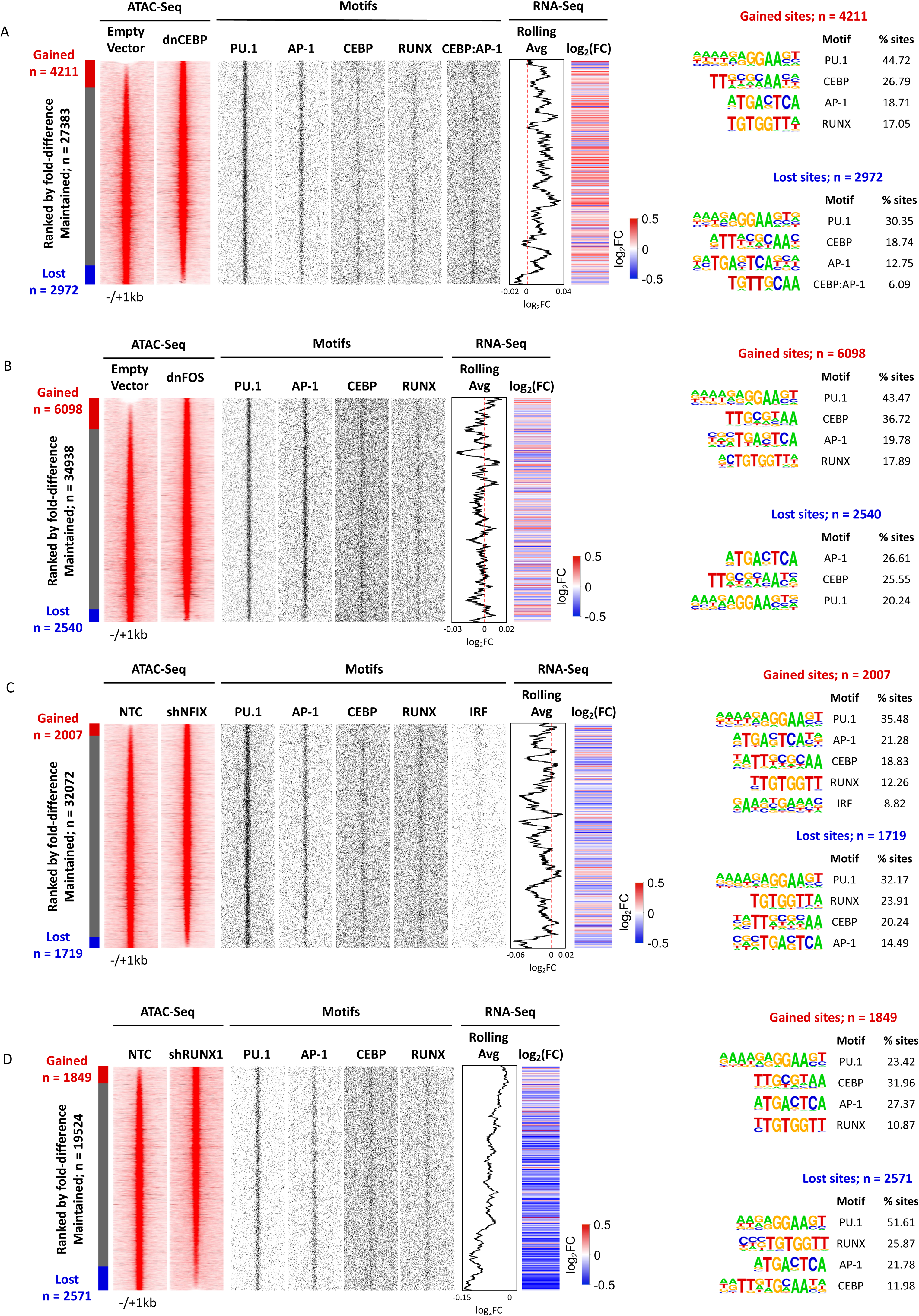
Perturbation of TF regulatory modules alters the chromatin landscape of FLT3-ITD primary AML cells. A-D: Density plot showing ATAC sites (red) changed after (A) dnCEBP, (B) dnFOS, (C) shNFIX or (D) shRUNX1 expression compared to an empty vector control (A,B) or shNTC (C,D). Data are ranked by normalized tag counts of ATAC peaks from control cells over peaks obtained from transduced cells. gained and lost open chromatin regions are indicated to the left of the bar. Middle panels: TF binding motifs (black) projected against hypersensitive sites are plotted alongside. Rolling average of gene expression values and fold change of gene expression are plotted alongside the DHS. Right panel: Enriched TF motifs in lost and gained open chromatin regions.

The analyses of gene expression after factor perturbation showed a complex regulatory relationship between the modules (Fig 5). All TF perturbations led to both an up- and down-regulation of genes. Inspection of genes that responded to NFIX and RUNX1 knock-down revealed that NFIX is in the same regulatory pathway as RUNX1, whereby *RUNX1* is strongly down-regulated after NFIX knockdown together with 56 other genes. RUNX1 depletion affected direct RUNX1 target loci driving macrophage differentiation such as *CSF1R* and *IRF8* ^19,20^ and multiple inflammatory genes. The indirect effects of RUNX1 down-regulation can also be seen in the NFIX knock-down although the fold change of many genes did not reach the significance level. Another cross-module response is seen after dnFOS expression which down-regulates NFIX and its homologue NFIA. It has been shown that AP-1 (JUN) and C/EBP family members can physically interact to drive macrophage differentiation ^21^ and our data show them to co-regulate a number of genes. In this context it is interesting to note that one the motifs enriched in open chromatin regions lost after dnC/EBP expression is a C/EBP/AP-1 composite motif (Fig 4a), suggesting that the two factors indeed cooperate.

**Figure 5.**
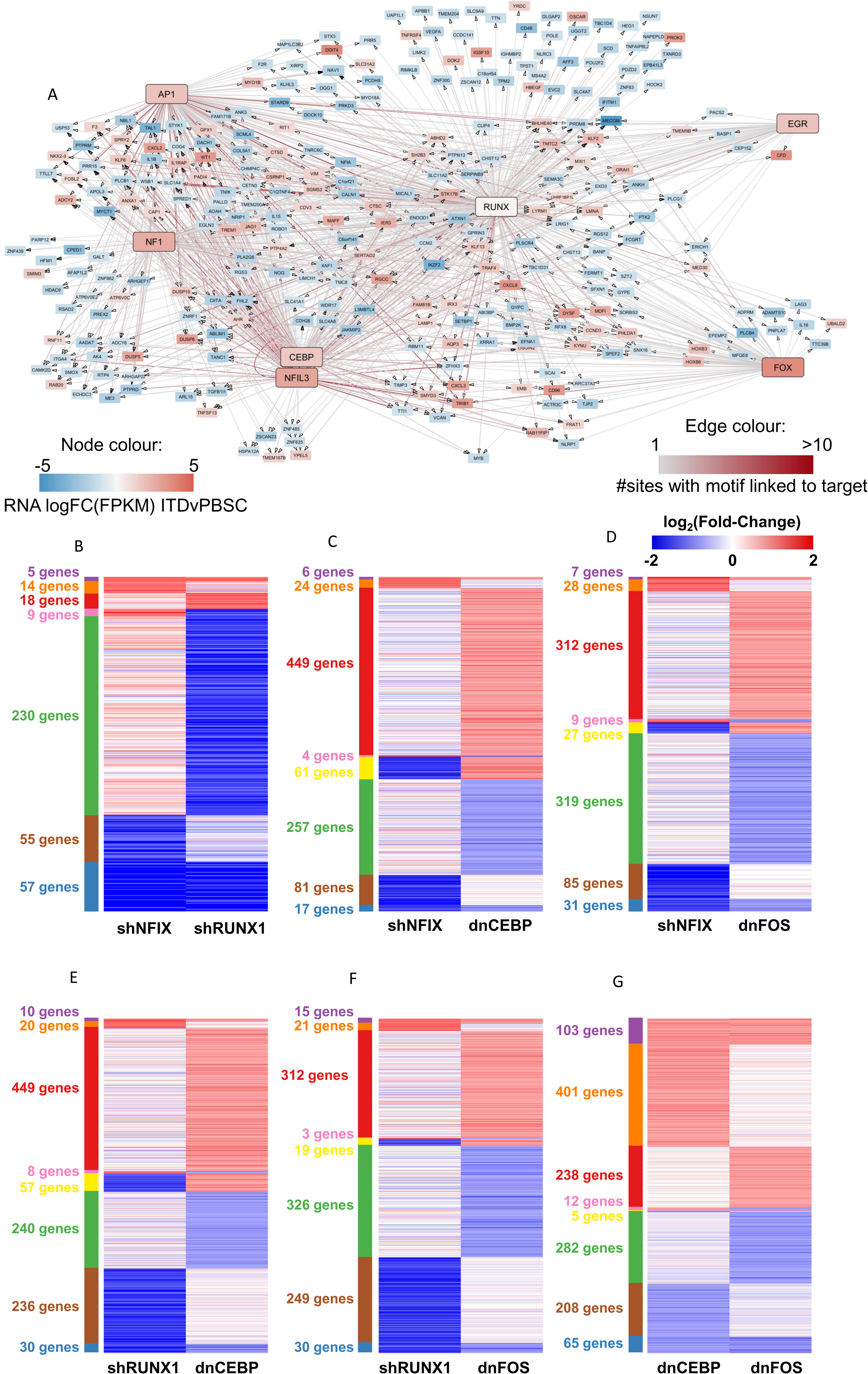
Perturbation assays reveal crosstalk between regulatory modules. A: Network showing the connection of the RUNX1 module to other indicated modules for genes upregulated (red) or down-regulated (blue) as compared to PBSCs. Node colour indicates the FC in RNA expression of the gene in FLT3-ITD AML compared to healthy PBSC. Edges indicate the presence of DHSs assigned to the gene containing motifs, edge colour corresponds to the number of interacting DHSs.B-G: Pairwise comparison of the log2FC of differentially expressed genes in primary FLT3-ITD AML cells (ITD-12) subjected to the indicated TF knock-downs compared to an empty vector control (dnCEBP, dnFOS) or shNTC (shRUNX1, shNFIX) respectively.

Taken together these few examples indicate an extensive crosstalk between the different regulatory modules. Perturbation of one module leads to a complex response of other modules which moves the GRN onto a new regulatory state.

### RUNX1 is an essential factor for the establishment of a FLT3-ITD-specific gene expression program regulating growth

Our experiments show that the perturbation of each selected TF module from the AML-specific GRN led to an abolition of AML colony forming ability and growth. To address the molecular basis of this finding, we concentrated on the RUNX1 module. Various AML sub-types, in particular core binding factor AMLs ^22–24^ are dependent on the presence of a wild-type copy of RUNX1. In addition, the analysis of leukemia reconstituting cells derived from induced pluripotent stem cells (iPSCs) from FLT3-ITD/NPM1 patients suggested an important role for this TF in this AML-sub-type as well ^25^. To ensure that our digital footprint-based module construction was valid, we validated binding sites by comparing them to previous ChIP-Seq experiments in FLT3-ITD/NPM1 patients and cell lines ^1^ (Fig S5B) and found that 79% of footprinted sites are also bound by RUNX1. In FLT3-ITD AML, the RUNX1 module strongly interconnects with the other modules (Fig 5A) to establish a FLT3-ITD-specific gene expression pattern (Fig S5A) and its motifs are the most enriched (note the co-enrichment of NFI motifs) (Fig S5B). The importance of RUNX1 in establishing this gene expression pattern is highlighted by the fact that in patients where both RUNX1 alleles are mutated, it is abolished (Figs S5 C – D) and FLT3-IITD specific genes are not expressed (Fig S5F). Moreover, as shown before ^1,2^ FLT3-ITD specifically expressed genes are highly enriched in the RUNX1 module (Fig S5I).

The fact that the depletion of aberrant or normal TF targets within AML specific GRNs affects AML, but not normal cells, gives rise to the hope that malignant epigenetic states can be reprogrammed, and cells can be driven into a cellular state that is incompatible with survival. TFs were thought to be “undruggable” but in recent years significant progress has been made to target these factors ^26^. We therefore probed the FLT3-ITD-specific GRN with the small molecule AI-10-91 (CBFβi) which disrupts the interaction between the RUNX DNA binding domain and its binding partner CBFβ ^27^. We first used a proximity ligation assay to determine the optimal time point to detect a complete dissociation of RUNX1::CBFβi (Fig 6A; Fig S6A) which we found to be 8 hours, suggesting that the complex is quite stable within the cell. ChIP experiments with primary cells treated with CBFβi confirmed a wide-spread loss of RUNX1 from the genome (Fig 6A, right panel). The inhibitor had a profound effect on the viability and colony forming ability of FLT3-ITD/NPM1 and FLT3-ITD AML cells but not healthy cells in culture (Fig 6B, Fig S6B). The CBFβ inhibitor efficiently killed the cells after 6 days. Furthermore, the cells from a patient with the double RUNX1 mutation (RUNX1(2x) did not respond to the inhibitor except when a very high dose was used, again demonstrating that inhibition operates via the RUNX1 module.

**Figure 6.**
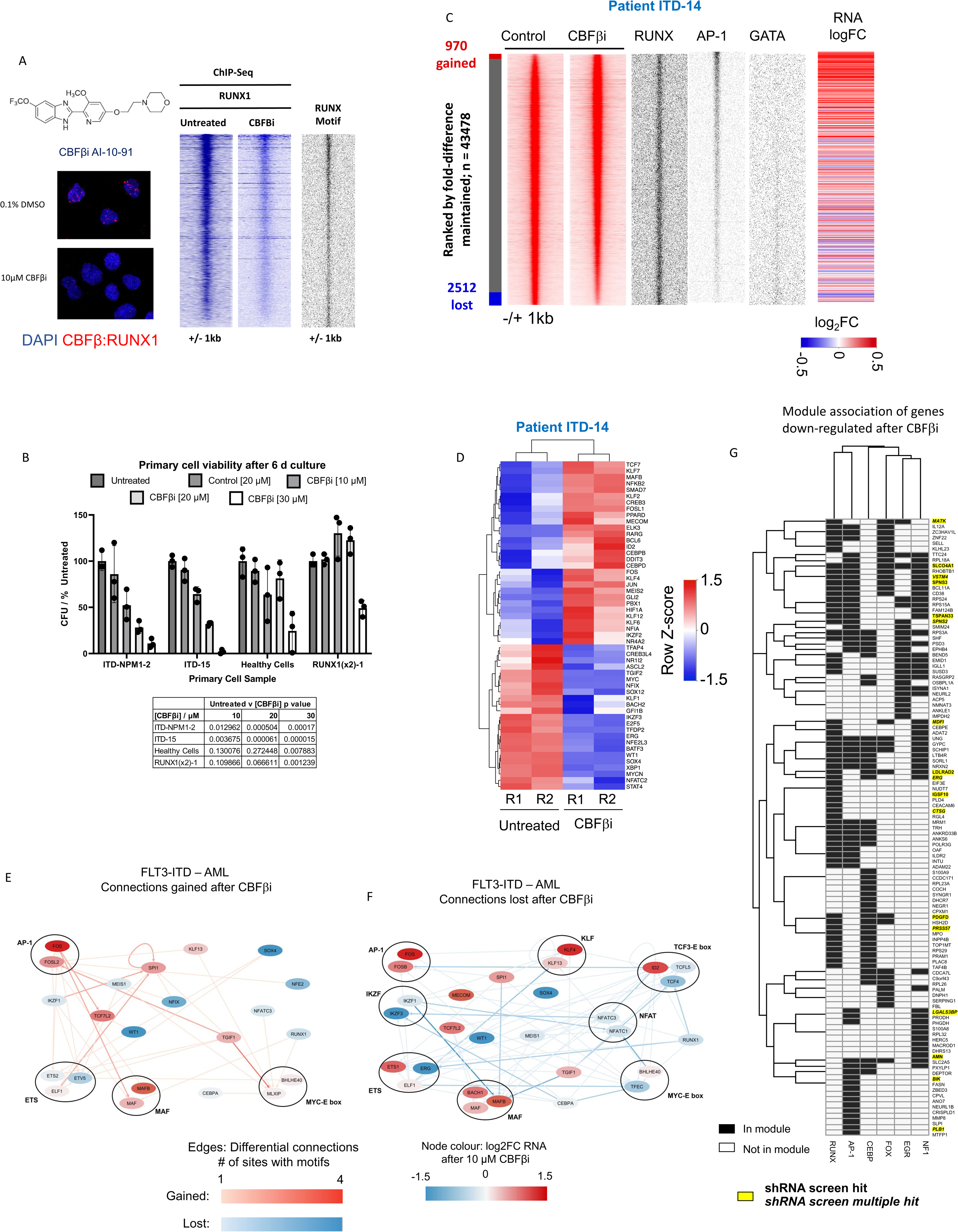
The RUNX1 regulatory module is essential for FLT3-ITD-specific gene expression - dissection of its contribution. A: Structure of CBFβi and proximity ligation assay of primary FLT3-ITD+ AML cells (ITD-14) with and without 10 μM CBFβI treatment. Right panel: Density plots of a ChIP experiments showing the genome wide signal of RUNX1 binding with and without CBFβi, as well as RUNX motifs present at the binding sites plotted alongside. B: Viability assays with the indicated primary cell types treated with increasing concentrations of CBFβΙ. RUNX1 (2x): cells from a patient with a double RUNX1 mutation. C: Density plot of ATAC-Seq analysis of primary FLT3-ITD patient cells (ITD-14) with and without 10 μM CBFβι ranked against each other according to fold-change with the indicated TF motifs at the open chromatin sites and the expression of the associated genes present plotted alongside. D: Unsupervised clustering of transcription factor gene expression of primary FLT3-ITD patient cells with and without CBFβi. E,F: Gained (E) and lost (F) connections in the FLT3-ITD-specific GRN before and after CBFβi treatment. Red edges show gained connections after CBFβI with blue showing those which are lost, node colour represents the fold change in RNA expression of the TF after CBFβi G: Genes downregulated in the RNA-seq data in 2 or more of the CBFβi treated patients. The heatmap shows the gene modules associated with each gene (black = associated, white = not associated). Yellow highlight: Genes scoring in our screen. Single hits in the screen in 1 or more samples are in written in bold, if there were multiple hits in 1 or more samples they are in italics. Genes not in the RUNX1, AP1, CEBP, EGR1, FOX, NF1 module are not included in this data. Hierarchical clustering was performed to group genes in similar modules.

Similar to the depletion of RUNX1 by shRNA mediated knock-down, the removal of RUNX1 from the genome led to not just a loss but also the gain of open chromatin regions and changes in gene expression (Fig 6C, Fig S6C, left panel). Again, inhibition yielded a complex response, with the expression of numerous FLT3-ITD/NPM1-specific genes in 3 different patients being up- or down regulated in similar patterns (Fig S6D) including multiple genes from the RUNX1 module (Fig S6D, right panel)). RUNX1 inhibition changed the expression of multiple transcription factor genes (Fig 6 D), suggesting that along-side the effect of CBFβi on cellular growth, that the GRN was rewired in response to treatment. We confirmed this idea using our ATAC-Seq data, and constructed GRNs from open chromatin sites that were gained and sites that were lost after inhibitor treatment (Fig 6E,F). The analysis of lost edges (Fig 6F) showed that multiple connections to the RUNX1 and to other TF nodes, such as the *CEBP* node, were lost, but other connections such as from and to the AP-1 family were gained (Fig 6E), indicating that not only connections to RUNX1, but also other TFs were altered. To close the circle, we therefore asked the question, how CBFβi treatment affected the genes belonging to the other 5 modules (NFI, AP-1, FOX, EGR, C/EBP) and how these associations were reflected in our shRNA screen (Fig 6G, Fig S6E). The analysis of genes down-regulated genes after treatment (Fig 6G) showed that the inhibition of RUNX1 activity had a profound influence on genes organised in the other modules, even when they were not overlapping with the RUNX1 modules. An example for this finding is *PLB1* which encodes for Phospholipase B1 whose overexpression has been observed in Glioblastoma and has been highlighted as a mRNA vaccine candidate (Fig 6G) ^28^. Down-regulated genes from other modules overlapping with the RUNX1 module scoring in our screen included MATK (Megakaryocyte-Associated Tyrosine Kinase) which is an important regulator of SRC kinases in blood cells ^29^. The analysis of up-regulated genes after CBFβi showed a strong enrichment of genes organised in the overlapping NFI, AP-1 and RUNX modules scoring in our screen. This included again *DUSP6* but also the genes encoding the Zn^++^ finger TFs KLF2 and KLF6 which have been shown to be repressed in AML cells and which are associated with myeloid differentiation ^30^.

In our final set of experiments, we asked the question of which cell type in primary AML cells and which genes were most affected by CBFβi. To this end, we cultured primary FLT3-ITD AML cells in the presence and absence of CBFβi and conducted single cell (sc) RNA-Seq experiments. Untreated cultures contained a mixture of early precursor cells, myeloid and erythroid cells of variable differentiation stages (Fig 7A. Fig S7A, see Fig S7B for a differentiation trajectory analysis). Inhibitor treatment reduced cell numbers with early precursors and myeloid cells being the most affected cell types (Fig 7D). The analysis of cell cycle-specifically expressed genes before and after inhibition showed a block in G1 which affected all cell types. The inspection of down-regulated genes in early progenitors revealed a strong down-regulation of cell cycle regulators and ribosomal genes (Fig 7 E), which was consistent with the cell cycle block. Also in this experiment, the cell cycle block was followed by cell death as measured by a viability dye assay.

**Figure 7.**
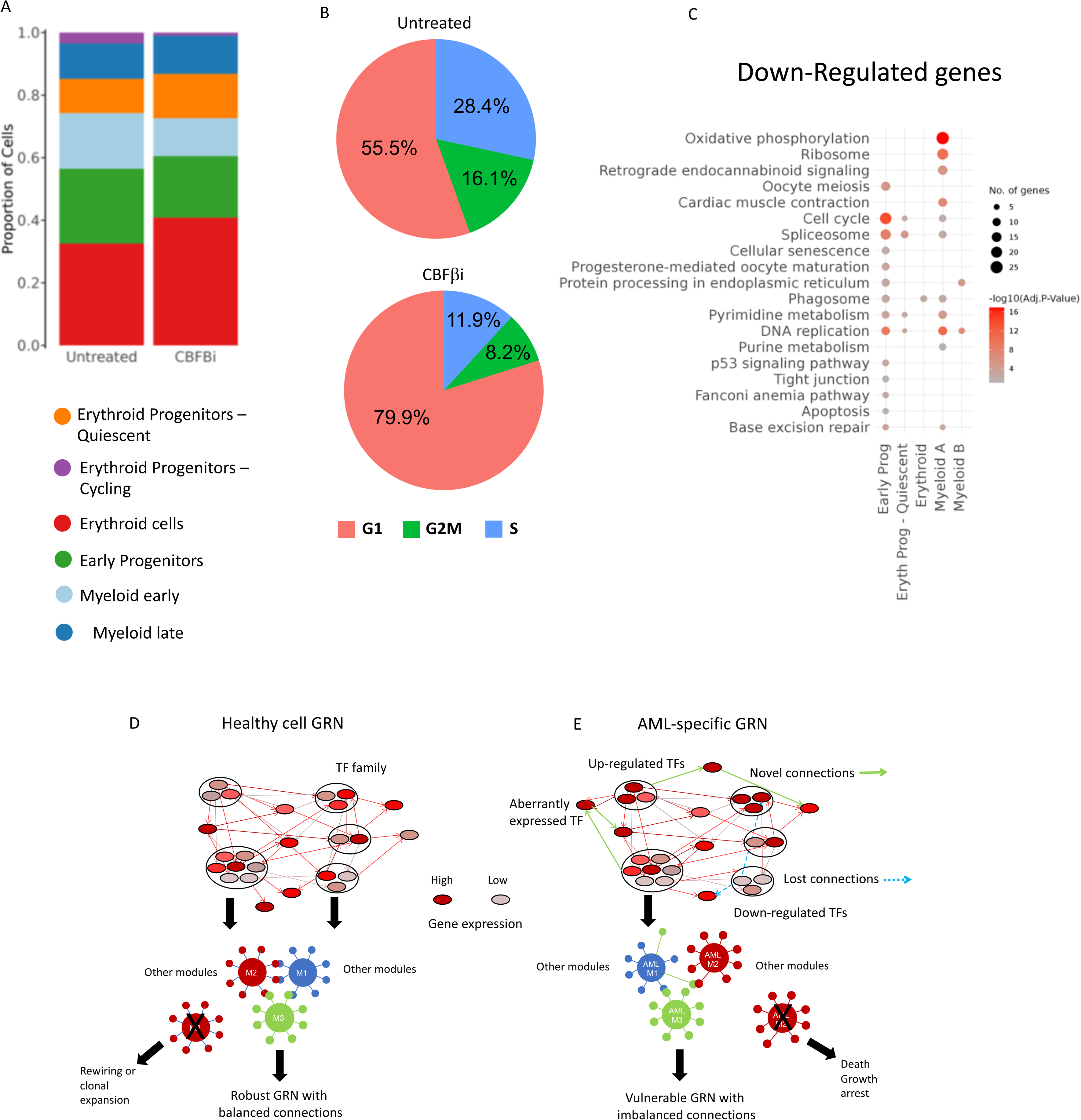
RUNX1 inactivation impedes cell cycle progression in blood progenitors. A: UMAP plot showing the proportion of the indicated cell types in the population with and without CBFbi treatment. B: Proportion of cells in the indicated cell cycle phase without (upper panel) and with (lower panel) CBFβi treatment. C: GO terms of genes down-regulated after CBFβi treatment. D, E: Model describing the difference between a robust healthy GRN in blood progenitors (D) and a vulnerable GRN found in malignant cells (E).

In summary, our experiments show that targeting an important TF comprising a highly connected node within an AML sub-type-specific GRN such as RUNX1 leads to a profound impact on other TF modules which rewires the GRN of AML cells, leading to a cell cycle block and eventually, cell death.

## Discussion

The present study shows how identification of an aberrant AML sub-type specific GRN based on the data collected by Assi et al., ^2^ leads to the identification of genes whose expression is vital for the maintenance of malignant cells. The hit-rate for our screen was very high, and we found numerous TF encoding genes which were important for the growth of AML, but not normal cells. Moreover, the identification of the down-stream targets of such TFs highlighted a complex web of interactions between genes which were essential for FLT3-ITD/NPM1 AML growth. This and our previous studies show that the idea of an AML sub-type specific GRN that maintains AML cells is not only true for groups affected by mutations of gene regulatory molecules themselves, but also for those with signalling mutations who generate chronic growth signals, such as the FLT3-ITD. Decades of clinical trials have tried to target aberrant signalling and have failed due to the development of drug resistance. Moreover, such treatments only affected fast growing cells, and not quiescent leukemic stem cells. Based on these data and those of others ^31,32^, we believe that the most viable way to eliminate malignant cells is to identify the factors crucial for tumour maintenance and reprogram cells into either normal cells that differentiate or change their GRNs into an unstable, non-viable state by identifying TFs that are essential for AML maintenance.

Our GRN analyses inform us as to which factors and genes to target. As schematically outlined in Fig 7D, in healthy cells, after hundreds of millions of years of evolution of complex multicellular organisms, the balance between self-renewal, differentiation and growth is highly robust and cells do not rely on one factor alone for differentiation, as exemplified by non-malignant clonal hematopoiesis. Moreover, as exemplified by the similarity of modules in the GRN shared between aberrant and normal cells (Fig 3D), all modules are in balance and all interact with each other (note the high cross module similarity), thus creating a stable network. With the possible exception of MLL-translocations which involve global activators of gene expression, it takes several mutations to derail normal hematopoietic differentiation. The majority of initiating mutations causing AML are found in members of the gene regulatory machinery. Differentiating mutant cells compensate for the loss or impediment of such activities by rewiring their gene regulatory networks, thus becoming dependent on alternative factors and pathways, which are often different to those of normal cells (Fig 7E). Mechanisms of this kind are often the upregulation of alternate signalling pathways, overexpression of TF genes or the ectopic activation of TF genes normally not expressed in these cells. This phenomenon can also be seen in the FLT3-ITD sub-type whose AML-specific GRN highlights various TF families which are associated with aberrant signalling (AP-1 family), are overexpressing (RUNX1) ^1,33^, or are aberrantly over-expressed (FOXC1, NFIX) ^2,16^ In addition, these sub-type specifically deregulated TFs form regulatory modules that contain multiple signalling genes that are up-regulated compared to healthy cells, such as the *DUSPs* or TNF-superfamily members (See Fig 5 A-F). Genes are regulated by a multitude of TFs which form interacting protein complexes on enhancers and promoters ^34,35^. Such interactions can be aberrant as exemplified by the formation of AML-specific protein complexes containing FOXC1 and RUNX1 ^36^

Recent experiments coupling degrons to oncoproteins or normal transcription factors have shown that only a small number of genes immediately respond to the degradation of one factor, and that it takes a number of hours until the whole GRN responds and changes the expression of multiple genes ^37^. It takes even longer until cells differentiate or die, and it is this feature which is relevant in a clinical setting. Our data highlight the molecular reason for this finding by showing that an AML-specific GRN sustaining leukemic growth is stabilised by multiple modules which cooperate in driving AML sub-type specific expression. The removal of one essential module leads to a rewiring of others, as exemplified by our detailed studies of the RUNX1 module. Here, the result is a profound cell cycle block followed by eventual cell death suggesting an inability of RUNX1-depleted cells to further rewire their GRN into a stable state and which could be the basis of therapy. A recent study applying a degron technology to target IKZF2 in MLL-driven AML ^38^ demonstrates that such a strategy is indeed feasible.

## Methods

A detailed description of the methods can be found in Supplementary Materials.

## Data and code availability

Data will be available on publication.

## Author contributions

D.J.L.C., R.L.-M., P.S.C., H.B., L.A., S.G.K., J.G., E.H., performed experiments. P.K and S.A.A. analysed data. J.B. provided essential reagents. O.H., P.N.C. and C.B. conceived and designed experiments and wrote the paper.

## Acknowledgements

We are grateful to the Genomics Birmingham Facility and to the University of Birmingham Flow Cytometry platform for their expert services. This work as funded by a grant from the Medical Research Council UK to C.B., O.H. and P.N.C (MR/S021469/1); by a grant from Blood Cancer UK to C.B. and P.N.C (15001) and by a grant from the National Institute of Health to J.B (R01 CA234478). D.J.L.C. is a John Goldman Fellow of Leukaemia UK (2021/JGF/001).

## References

1. Cauchy, P., James, S.R., Zacarias-Cabeza, J., Ptasinska, A., Imperato, M.R., Assi, S.A., Piper, J., Canestraro, M., Hoogenkamp, M., Raghavan, M., et al. (2015). Chronic FLT3-ITD Signaling in Acute Myeloid Leukemia Is Connected to a Specific Chromatin Signature. Cell Rep 12, 821–836. 10.1016/j.celrep.2015.06.069.

2. Assi, S.A., Imperato, M.R., Coleman, D.J.L., Pickin, A., Potluri, S., Ptasinska, A., Chin, P.S., Blair, H., Cauchy, P., James, S.R., et al. (2019). Subtype-specific regulatory network rewiring in acute myeloid leukemia. Nat Genet 51, 151–162. 10.1038/s41588-018-0270-1.

3. Stirewalt, D.L., and Radich, J.P. (2003). The role of FLT3 in haematopoietic malignancies. Nat Rev Cancer 3, 650–665. 10.1038/nrc1169.

4. Corces-Zimmerman, M.R., Hong, W.J., Weissman, I.L., Medeiros, B.C., and Majeti, R. (2014). Preleukemic mutations in human acute myeloid leukemia affect epigenetic regulators and persist in remission. Proceedings of the National Academy of Sciences of the United States of America 111, 2548–2553. 10.1073/pnas.1324297111.

5. Miles, L.A., Bowman, R.L., Merlinsky, T.R., Csete, I.S., Ooi, A.T., Durruthy-Durruthy, R., Bowman, M., Famulare, C., Patel, M.A., Mendez, P., et al. (2020). Single-cell mutation analysis of clonal evolution in myeloid malignancies. Nature 587, 477–482. 10.1038/s41586-020-2864-x.

6. Daver, N., Schlenk, R.F., Russell, N.H., and Levis, M.J. (2019). Targeting FLT3 mutations in AML: review of current knowledge and evidence. Leukemia 33, 299–312. 10.1038/s41375-018-0357-9.

7. McMahon, C.M., and Perl, A.E. (2019). Management of primary refractory acute myeloid leukemia in the era of targeted therapies. Leuk Lymphoma 60, 583–597. 10.1080/10428194.2018.1504937.

8. Gallipoli, P., Giotopoulos, G., Tzelepis, K., Costa, A.S.H., Vohra, S., Medina-Perez, P., Basheer, F., Marando, L., Di Lisio, L., Dias, J.M.L., et al. (2018). Glutaminolysis is a metabolic dependency in FLT3(ITD) acute myeloid leukemia unmasked by FLT3 tyrosine kinase inhibition. Blood 131, 1639–1653. 10.1182/blood-2017-12-820035.

9. Choi, A., Jang, I., Han, H., Kim, M.S., Choi, J., Lee, J., Cho, S.Y., Jun, Y., Lee, C., Kim, J., et al. (2021). iCSDB: an integrated database of CRISPR screens. Nucleic Acids Res 49, D956–D961. 10.1093/nar/gkaa989.

10. Dekker, J., Rippe, K., Dekker, M., and Kleckner, N. (2002). Capturing chromosome conformation. Science 295, 1306–1311. 10.1126/science.1067799.

11. Piper, J., Elze, M.C., Cauchy, P., Cockerill, P.N., Bonifer, C., and Ott, S. (2013). Wellington: a novel method for the accurate identification of digital genomic footprints from DNase-seq data. Nucleic Acids Res 41, e201. 10.1093/nar/gkt850.

12. Gale, R.E., Green, C., Allen, C., Mead, A.J., Burnett, A.K., Hills, R.K., Linch, D.C., and Medical Research Council Adult Leukaemia Working, P. (2008). The impact of FLT3 internal tandem duplication mutant level, number, size, and interaction with NPM1 mutations in a large cohort of young adult patients with acute myeloid leukemia. Blood 111, 2776–2784. 10.1182/blood-2007-08-109090.

13. Cancer Genome Atlas Research, N. (2013). Genomic and epigenomic landscapes of adult de novo acute myeloid leukemia. The New England journal of medicine 368, 2059–2074. 10.1056/NEJMoa1301689.

14. Martinez-Soria, N., McKenzie, L., Draper, J., Ptasinska, A., Issa, H., Potluri, S., Blair, H.J., Pickin, A., Isa, A., Chin, P.S., et al. (2018). The Oncogenic Transcription Factor RUNX1/ETO Corrupts Cell Cycle Regulation to Drive Leukemic Transformation. Cancer Cell 34, 626–642 e628. 10.1016/j.ccell.2018.08.015.

15. Olive, M., Williams, S.C., Dezan, C., Johnson, P.F., and Vinson, C. (1996). Design of a C/EBP-specific, dominant-negative bZIP protein with both inhibitory and gain-of-function properties. J Biol Chem 271, 2040–2047. 10.1074/jbc.271.4.2040.

16. Somerville, T.D., Wiseman, D.H., Spencer, G.J., Huang, X., Lynch, J.T., Leong, H.S., Williams, E.L., Cheesman, E., and Somervaille, T.C. (2015). Frequent Derepression of the Mesenchymal Transcription Factor Gene FOXC1 in Acute Myeloid Leukemia. Cancer Cell 28, 329–342. 10.1016/j.ccell.2015.07.017.

17. O’Connor, C., Campos, J., Osinski, J.M., Gronostajski, R.M., Michie, A.M., and Keeshan, K. (2015). Nfix expression critically modulates early B lymphopoiesis and myelopoiesis. PLoS One 10, e0120102. 10.1371/journal.pone.0120102.

18. Hall, T., Walker, M., Ganuza, M., Holmfeldt, P., Bordas, M., Kang, G., Bi, W., Palmer, L.E., Finkelstein, D., and McKinney-Freeman, S. (2018). Nfix Promotes Survival of Immature Hematopoietic Cells via Regulation of c-Mpl. Stem Cells 36, 943–950. 10.1002/stem.2800.

19. Sauter, K.A., Bouhlel, M.A., O’Neal, J., Sester, D.P., Tagoh, H., Ingram, R.M., Pridans, C., Bonifer, C., and Hume, D.A. (2013). The function of the conserved regulatory element within the second intron of the mammalian Csf1r locus. PLoS One 8, e54935. 10.1371/journal.pone.0054935.

20. Holtschke, T., Lohler, J., Kanno, Y., Fehr, T., Giese, N., Rosenbauer, F., Lou, J., Knobeloch, K.P., Gabriele, L., Waring, J.F., et al. (1996). Immunodeficiency and chronic myelogenous leukemia-like syndrome in mice with a targeted mutation of the ICSBP gene. Cell 87, 307–317. 10.1016/s0092-8674(00)81348-3.

21. Cai, D.H., Wang, D., Keefer, J., Yeamans, C., Hensley, K., and Friedman, A.D. (2008). C/EBP alpha:AP-1 leucine zipper heterodimers bind novel DNA elements, activate the PU.1 promoter and direct monocyte lineage commitment more potently than C/EBP alpha homodimers or AP-1. Oncogene 27, 2772–2779. 10.1038/sj.onc.1210940.

22. Ben-Ami, O., Friedman, D., Leshkowitz, D., Goldenberg, D., Orlovsky, K., Pencovich, N., Lotem, J., Tanay, A., and Groner, Y. (2013). Addiction of t(8;21) and inv(16) acute myeloid leukemia to native RUNX1. Cell reports 4, 1131–1143. 10.1016/j.celrep.2013.08.020.

23. Loke, J., Assi, S.A., Imperato, M.R., Ptasinska, A., Cauchy, P., Grabovska, Y., Soria, N.M., Raghavan, M., Delwel, H.R., Cockerill, P.N., et al. (2017). RUNX1-ETO and RUNX1-EVI1 Differentially Reprogram the Chromatin Landscape in t(8;21) and t(3;21) AML. Cell reports 19, 1654–1668. 10.1016/j.celrep.2017.05.005.

24. Goyama, S., Schibler, J., Cunningham, L., Zhang, Y., Rao, Y., Nishimoto, N., Nakagawa, M., Olsson, A., Wunderlich, M., Link, K.A., et al. (2013). Transcription factor RUNX1 promotes survival of acute myeloid leukemia cells. J Clin Invest 123, 3876–3888. 10.1172/JCI68557.

25. Wesely, J., Kotini, A.G., Izzo, F., Luo, H., Yuan, H., Sun, J., Georgomanoli, M., Zviran, A., Deslauriers, A.G., Dusaj, N., et al. (2020). Acute Myeloid Leukemia iPSCs Reveal a Role for RUNX1 in the Maintenance of Human Leukemia Stem Cells. Cell reports 31, 107688. 10.1016/j.celrep.2020.107688.

26. Bushweller, J.H. (2019). Targeting transcription factors in cancer - from undruggable to reality. Nat Rev Cancer 19, 611–624. 10.1038/s41568-019-0196-7.

27. Illendula, A., Gilmour, J., Grembecka, J., Tirumala, V.S.S., Boulton, A., Kuntimaddi, A., Schmidt, C., Wang, L., Pulikkan, J.A., Zong, H., et al. (2016). Small Molecule Inhibitor of CBFbeta-RUNX Binding for RUNX Transcription Factor Driven Cancers. EBioMedicine 8, 117–131. 10.1016/j.ebiom.2016.04.032.

28. Lin, H., Wang, K., Xiong, Y., Zhou, L., Yang, Y., Chen, S., Xu, P., Zhou, Y., Mao, R., Lv, G., et al. (2022). Identification of Tumor Antigens and Immune Subtypes of Glioblastoma for mRNA Vaccine Development. Front Immunol 13, 773264. 10.3389/fimmu.2022.773264.

29. Lee, B.C., Avraham, S., Imamoto, A., and Avraham, H.K. (2006). Identification of the nonreceptor tyrosine kinase MATK/CHK as an essential regulator of immune cells using Matk/CHK-deficient mice. Blood 108, 904–907. 10.1182/blood-2005-12-4885.

30. Humbert, M., Halter, V., Shan, D., Laedrach, J., Leibundgut, E.O., Baerlocher, G.M., Tobler, A., Fey, M.F., and Tschan, M.P. (2011). Deregulated expression of Kruppel-like factors in acute myeloid leukemia. Leuk Res 35, 909–913. 10.1016/j.leukres.2011.03.010.

31. Papapetrou, E.P., and Lee, D.F. (2021). Reprogramming and cancer. Stem Cell Res 52, 102249. 10.1016/j.scr.2021.102249.

32. Bonifer, C., and Cockerill, P.N. (2015). Chromatin Structure Profiling Identifies Crucial Regulators of Tumor Maintenance. Trends in cancer 1, 157–160. 10.1016/j.trecan.2015.10.003.

33. Behrens, K., Maul, K., Tekin, N., Kriebitzsch, N., Indenbirken, D., Prassolov, V., Muller, U., Serve, H., Cammenga, J., and Stocking, C. (2017). RUNX1 cooperates with FLT3-ITD to induce leukemia. J Exp Med 214, 737–752. 10.1084/jem.20160927.

34. Hafner, A., and Boettiger, A. (2023). The spatial organization of transcriptional control. Nat Rev Genet 24, 53–68 10.1038/s41576-022-00526-0.

35. Lim, B., and Levine, M.S. (2021). Enhancer-promoter communication: hubs or loops? Curr Opin Genet Dev 67, 5–9. 10.1016/j.gde.2020.10.001.

36. Simeoni, F., and Somervaille, T.C. (2021). Enhancer recruitment of a RUNX1, HDAC1 and TLE3 co-repressor complex by mis-expressed FOXC1 blocks differentiation in acute myeloid leukemia. Mol Cell Oncol 8, 2003161. 10.1080/23723556.2021.2003161.

37. Stengel, K.R., Ellis, J.D., Spielman, C.L., Bomber, M.L., and Hiebert, S.W. (2021). Definition of a small core transcriptional circuit regulated by AML1-ETO. Mol Cell 81, 530–545 e535. 10.1016/j.molcel.2020.12.005.

38. Park, S.M., Miyamoto, D.K., Han, G.Y.Q., Chan, M., Curnutt, N.M., Tran, N.L., Velleca, A., Kim, J.H., Schurer, A., Chang, K., et al. (2023). Dual IKZF2 and CK1alpha degrader targets acute myeloid leukemia cells. Cancer Cell 41, 726–739 e711. 10.1016/j.ccell.2023.02.010.

