## Supplemental Materials for "Gene regulatory network analysis predicts cooperating transcription factor regulons required for FLT3-ITD+ AML growth"

### Supplementary Materials

#### List of Supplementary Materials

#### 1. Supplementary Table 1: Patient Data

#### 2. Supplementary Figures and Figure legends

#### 3. Supplementary Materials and Methods

#### 4. Supplementary References

**Supplementary Table S1:**

**Patient Data**

| Patient Code | Mutations | Sex | wbc | Case |
|---|---|---|---|---|
| ITD-12 | 46XX, 7q-, i(13), DNMT3A, U2AF1 and FLT3-ITD | F | 65 | Presentation |
| ITD-13 | 46XY, FLT3-ITD, JAK2, CEBPA, | M | 376 | Presentation |
| ITD-14 | 46XX, FLT3-ITD, NPM1, ANKRD26, CEBPA (10%), ESXH2, TET2x2, ZRSR2 | F | 300 | Presentation |
| ITD-15 | 46 XY, FLT3 ITD, CHEK2, CUX1 | M | 17 | Relapse |
| ITD-NPM1-2 | 46XX, FLT3-ITD, NPM1 | F | 7 | Relapse |
| ITD-NPM1-6 | 46XX, FLT3-ITD, NPM1, WT1, DNMT3A, TET2, PHF6 | F | 195 | Presentation |
| RUNX1(x2)-1 | 46XY del 21q, RUNX1x2, TET2, STAG2, BCOR, IKZF1, EZH2 | M | 78 | Presentation after MDS |

Information on patient samples used in this paper. For other patients used in analysis see Assi et al 2019[1].

Figure S1

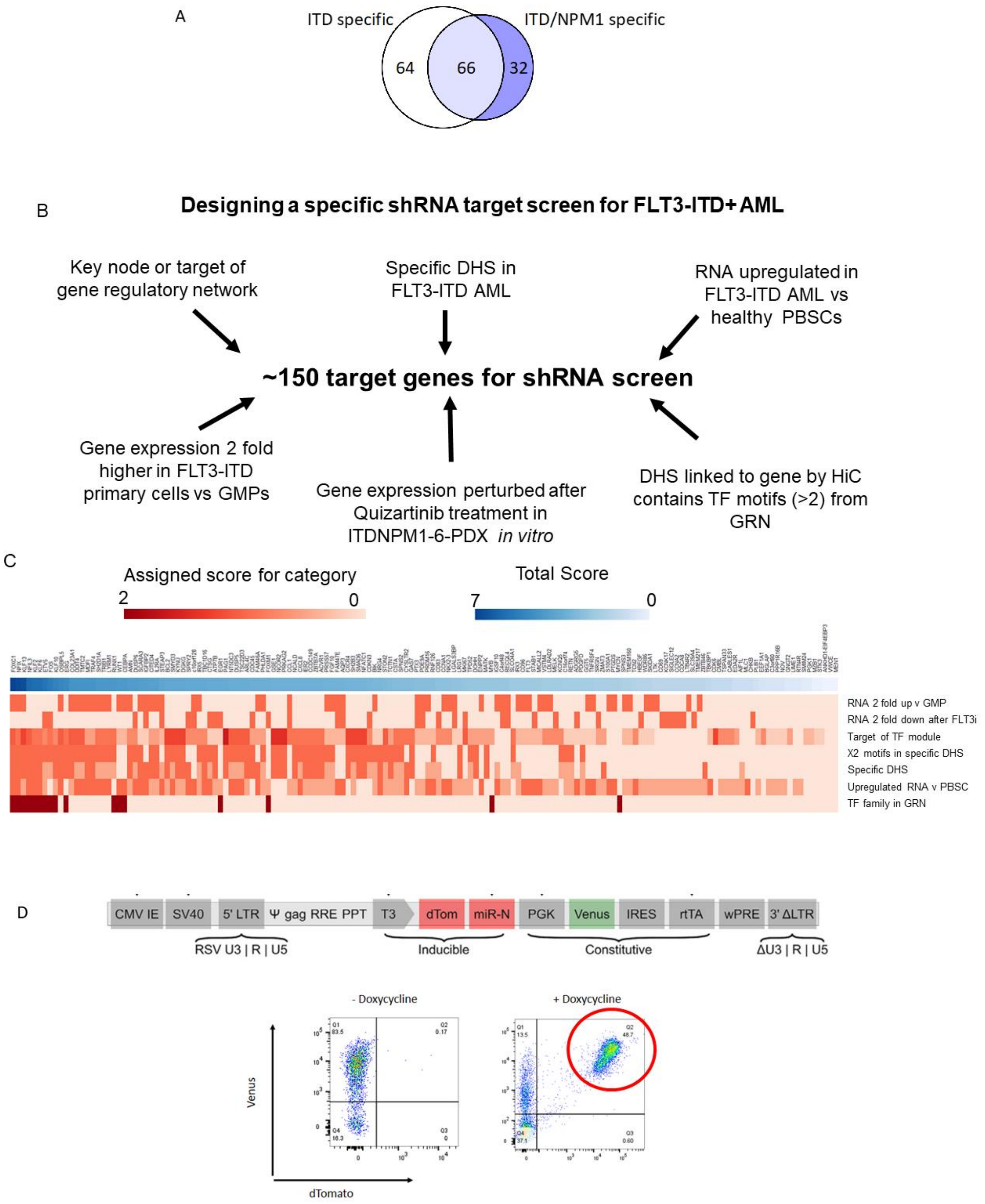


### Figure S1 (related to Figures 1&2). Design of shRNA screen targeting the FLT3-ITD GRN

A: Overlap of shared edges in FLT3-ITD and FLT3-ITD/NPM1 GRNs. B: Target genes for the shRNA screen were scored based on multiomic data from Assi et al 2019[1] and genes perturbed after treatment of ITD-NPM1-6-PDX primary cells treated with 10 nM quizartinib in vitro. C: Heatmap of scoring of genes selected for the FLT3-ITD screen based upon the criteria shown in B. Red heatmap shows the score of the gene for each category with the blue bar showing the total score. D: Structure of shRNA expressing lentivirus (pL40c) and marker gene expression after library transduction with and without Doxycycline treatment.

Figure S2

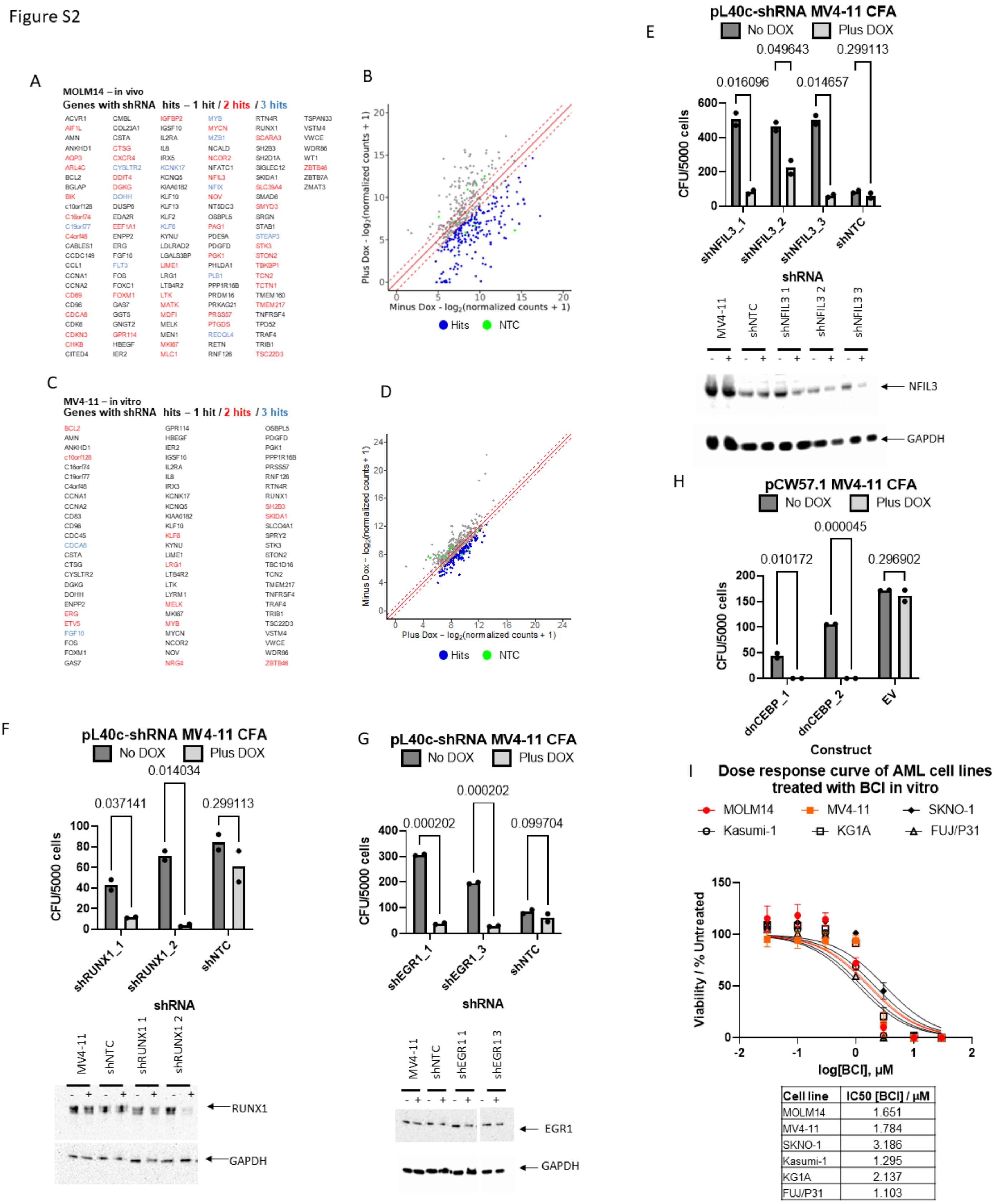


**Figure S2 (Related to Figure 2): Screening results and manual validation of selected hits in FLT3-ITD cell lines.**

A: MOLM14 in vivo screen results showing genes with shRNAs lost in the population, genes with multiple shRNA hits are highlighted. B: Scatter plot showing log2 shRNA frequency in MOLM14 in vivo screen, with lost shRNAs highlighted. C: MV4-11 in vitro screen results showing genes with shRNAs lost in the population, genes with multiple shRNA hits are highlighted. D: Scatter plot showing log2 shRNA frequency in MV4-11 in vitro screen, with lost shRNAs highlighted. E-H validation of GRN targets in MV4-11 cell lines, performed in duplicate for each construct, p-values were calculated using Student's t-test. Constructs that did not show a decrease in protein expression after 72 h induction were excluded from the analysis. E: Colony formation assays of MV4-11 cells transduced with shRNA targeting NFIL3. Induction of the shRNA knockdown of NFIL3 by doxycycline caused a decrease in colony formation compared to uninduced cells. The Western blot below shows a decrease in NFIL3 protein expression after 72 h induction, with GAPDH included as a control. F: Colony formation assays of MV4-11 cells transduced with shRNA targeting RUNX1. Induction of the shRNA knockdown of RUNX1 by doxycycline caused a decrease in colony formation compared to uninduced cells. The Western blot below shows a decrease in RUNX1 protein expression after 72 h induction, with GAPDH included as a control. G: Colony formation assays of MV4-11 cells transduced with shRNA targeting EGR1. Induction of the shRNA knockdown of EGR1 by doxycycline caused a decrease in colony formation compared to uninduced cells. The Western blot below shows a decrease in EGR1 protein expression after 72 h induction, with GAPDH included as a control. H: Expression of a dominant negative CEBP in MV4-11 clones showed a decrease in colony forming ability after induction compared to uninduced cells and MV4-11s transduced with an empty vector control. I: Dose response curves of cell lines treated with DUSP1/6 inhibitor BCI. FLT3-ITD cell lines show sensitivity to the inhibitor at a similar level to cell lines with MAPK activating mutations (Kasumi-1, P31/FUJ) whilst those without show marginally decreased sensitivity, although all AML cell lines respond to the inhibitor. Means calculated from n=3 are plotted with ± SEM and IC50 are shown below.

Figure S3

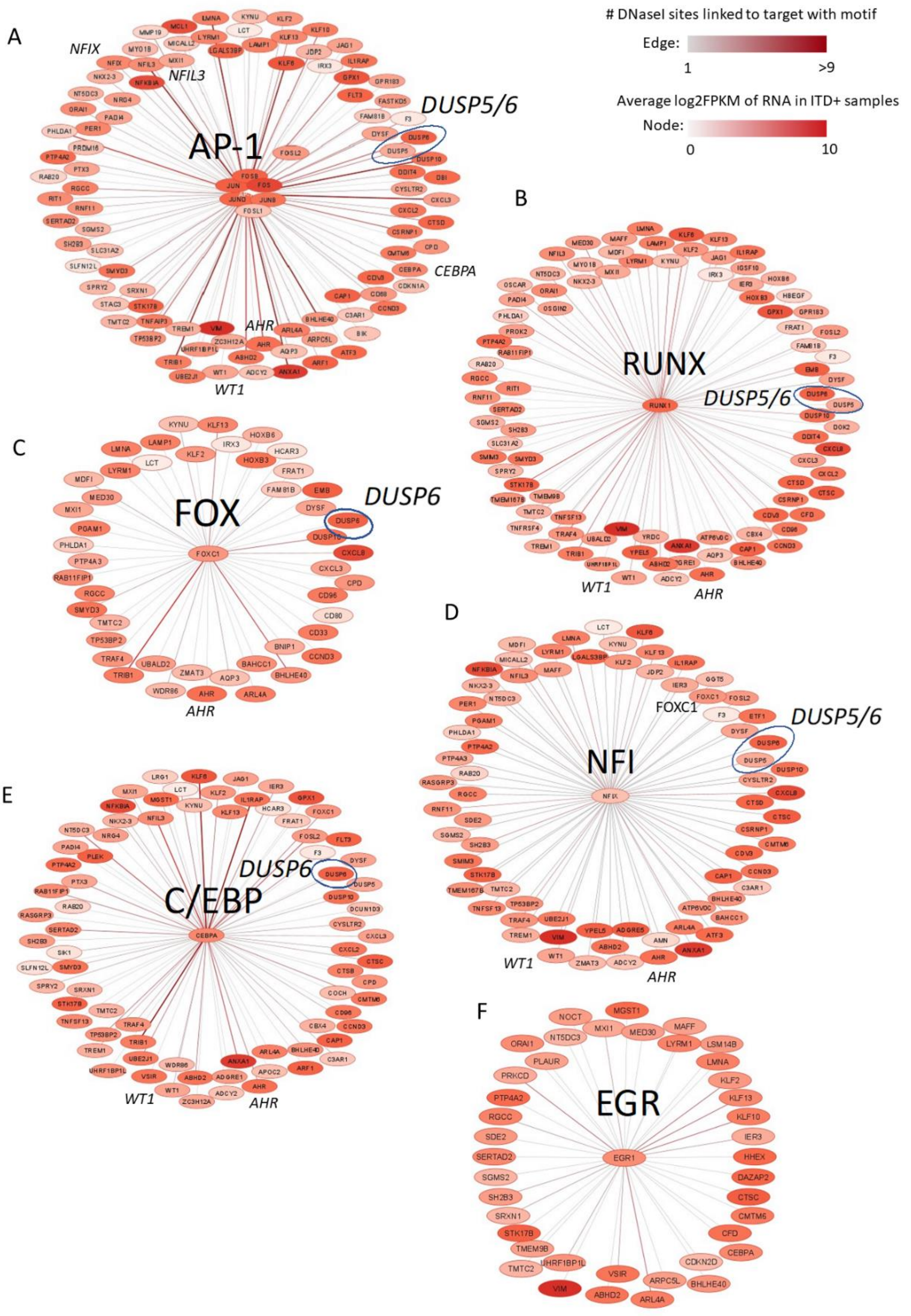


**Figure S3 (related to figure 3) TF modules of upregulated genes in FLT3-ITD AML:** A-F: AP-1, FOX, RUNX, NFI, C/EBP and EGR regulatory modules of FLT3-ITD AML specifically expressed genes as compared to PBSCs. Node colour indicates gene expression in FLT3-ITD+ AML samples (FPKM). Edges indicate an interaction between TF family and target genes, with the colour of the edge denoting the number of regions of open chromatin containing the TF family motif linked to the target gene by HiC or nearest gene (within 200 kb). TFs that are associated with more than one module are highlighted.

Figure S4

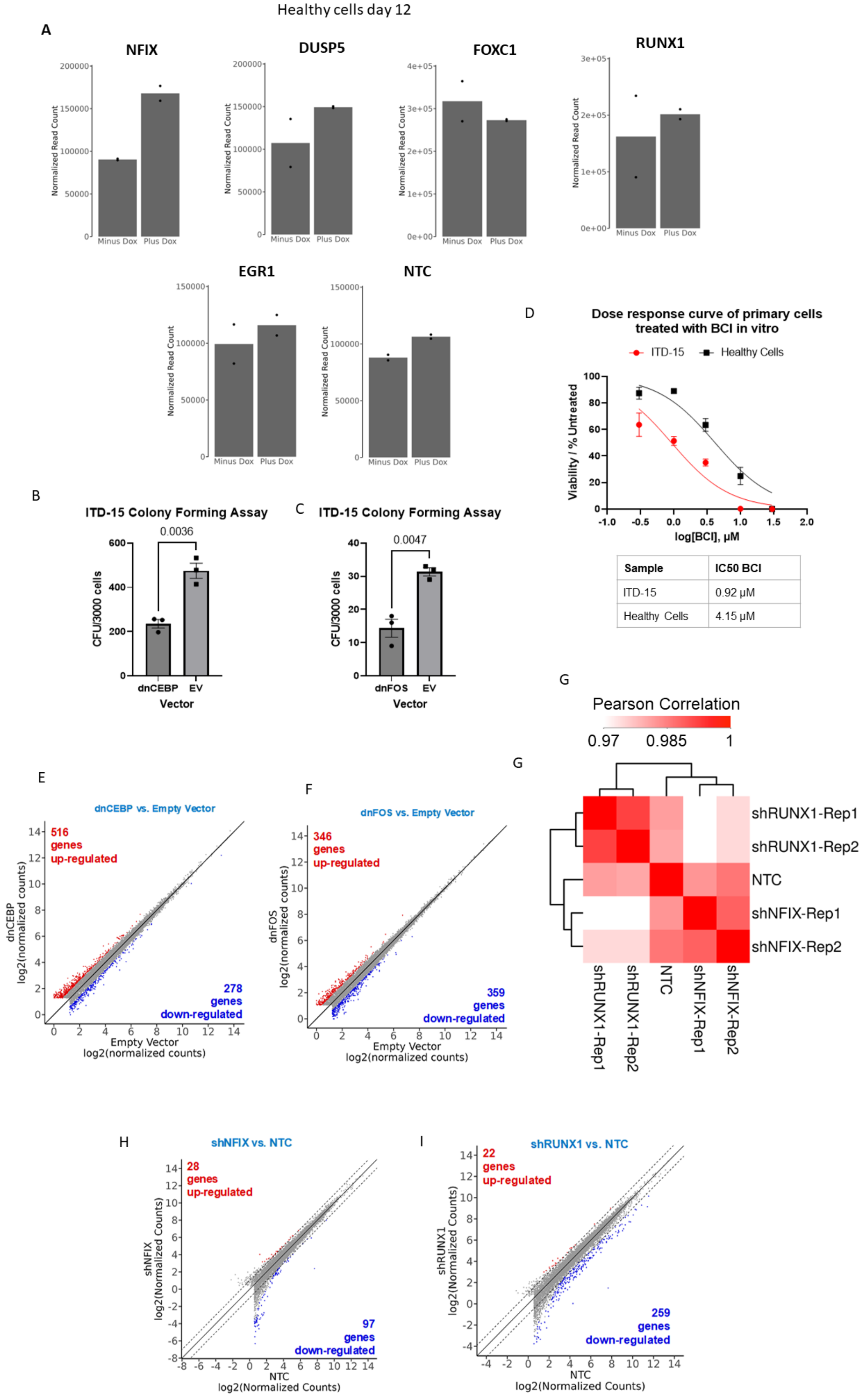
Healthy cells day 12
A
NFIX
DUSP5
FOXC1
RUNX1
EGR1
NTC
Normalized Read Count
Minus Dox
Plus Dox
B
ITD-15 Colony Forming Assay
0.0036
CFU/3000 cells
dnCEBP
EV
Vector
C
ITD-15 Colony Forming Assay
0.0047
dnFOS
D
Dose response curve of primary cells treated with BCI in vitro
ITD-15
Healthy Cells
Viability / % Untreated
log[BCI], μM
Sample
IC50 BCI
ITD-15
0.92 μM
Healthy Cells
4.15 μM
G
Pearson Correlation
0.97
0.985
1
shRUNX1-Rep1
shRUNX1-Rep2
NTC
shNFIX-Rep1
shNFIX-Rep2
E
dnCEBP vs. Empty Vector
516 genes up-regulated
278 genes down-regulated
dnCEBP log2(normalized counts)
Empty Vector log2(normalized counts)
F
dnFOS vs. Empty Vector
346 genes up-regulated
359 genes down-regulated
dnFOS log2(normalized counts)
H
shNFIX vs. NTC
28 genes up-regulated
97 genes down-regulated
shNFIX log2(Normalized Counts)
NTC log2(Normalized Counts)
I
shRUNX1 vs. NTC
22 genes up-regulated
259 genes down-regulated
shRUNX1 log2(Normalized Counts)

**Figure S4 (related to Figure 4). TF perturbation experiments in primary cells**

A: Bar charts showing the results of shRNA screen in healthy cells focused on 6 targets. DNA was harvested and libraries were prepared after 12 days of culture with or without doxycycline induction (n=2). B, C: Bar charts showing the effect of lentivirally transduced dominant negative CEBP (B) and FOS (C) peptide on colony forming ability of FLT3-ITD primary AML cells with and without Doxycycline stimulation. Experiments were performed in triplicate, mean colony formation per 5000 cells seeded are plotted and p values have been calculated by Student's t-test. Individual data points are plotted with the bar showing mean colony formation ± SEM. D: Dose response curve of treatment of FLT3-ITD primary AML cells and a healthy control with different concentrations of the DUSP inhibitor BCI as indicated. Means calculated from n=3 are plotted with ± SEM and IC50 are shown below. E,F: Scatter plot of mRNA expression patterns of FLT3-ITD primary AML cells with and without induction of dnCEBP and dnFOS compared to an empty vector control. H, I: Scatter plot of gene expression patterns of FLT3-ITD primary AML cells with and without expression of the indicated shRNAs compared to a non-targeting control. G: Pearson correlation between replicates of shRNA knockdown RNA-seq.

Figure S5

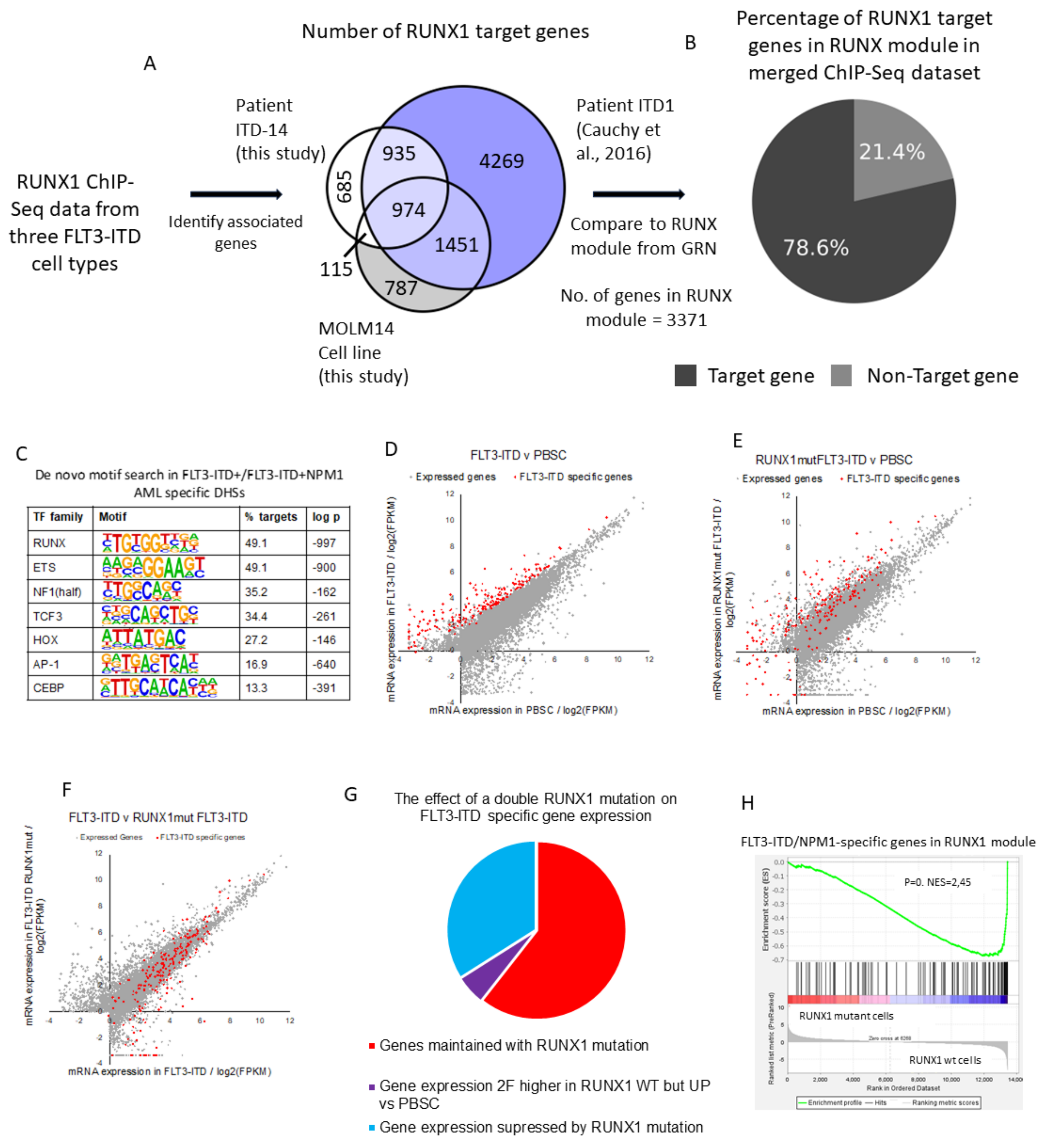


**Figure S5 (related to Figure 5) RUNX1 plays a key role in maintenance of the FLT3-ITD AML phenotype:**

A: Venn diagrams showing integration of three RUNX1 ChIP data-sets from primary FLT3-ITD AML cells and cell lines. B: Pie chart showing the percentage of RUNX1 target genes in RUNX module in merged RUNX1 ChIP-Seq dataset.

C: Table of enriched TF motifs in FLT3-ITD AML specific open chromatin regions compared to healthy PBSCs (Assi et al., 2019). D: Scatter plot comparing log2(FPKM) gene expression in FLT3-ITD AML to PBSCs. Genes specifically upregulated in ITD+ AML (>2 fold change, p<0.1) are highlighted in red. E: Scatter plot comparing log2(FPKM) gene expression in FLT3-ITD AML with a double RUNX1 mutation to PBSCs. Genes specifically upregulated in ITD+ AML (>2 fold change, p<0.1) are highlighted in red. F: Scatter plot comparing log2(FPKM) gene expression in FLT3-ITD AML to FLT3-ITD AML with a double RUNX1 mutation. Genes specifically upregulated in ITD+ AML (>2 fold change, p<0.1) are highlighted in red. G: The effect of a double RUNX1 mutation on FLT3-ITD specific gene expression in AML samples. H: GSEA showing the distribution of expression of FLT3-ITD specific genes in the RUNX1 module between AML samples with FLT3-ITD and RUNX1 WT or FLT3-ITD RUNX1 mutant cells.

Figure S6

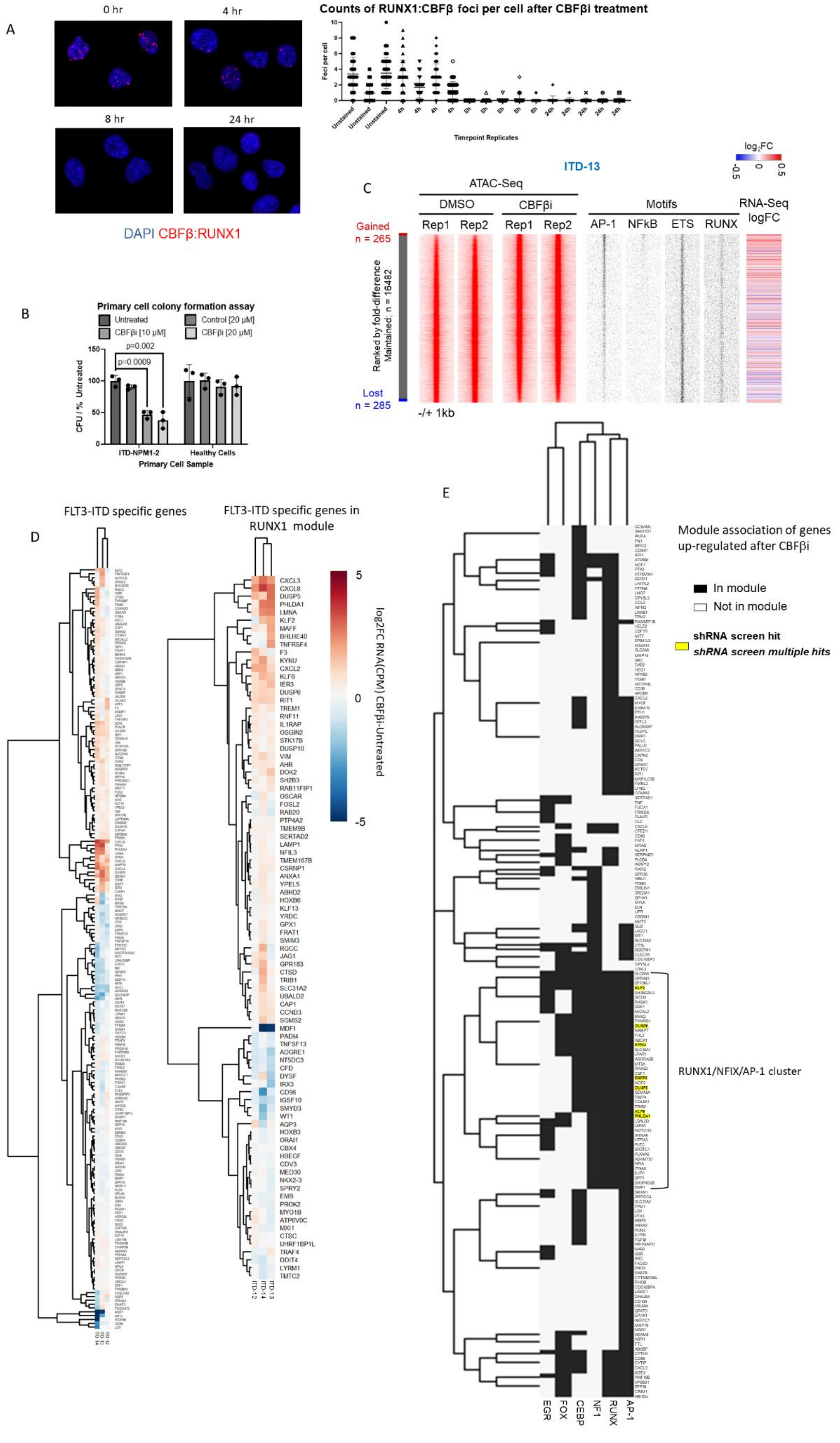

A
0 hr
4 hr
8 hr
24 hr
DAPI CBFβ:RUNX1
Counts of RUNX1:CBFβ foci per cell after CBFβi treatment
Foci per cell
Timepoint Replicates
B
Primary cell colony formation assay
Untreated
Control [20 μM]
CBFβi [10 μM]
CBFβi [20 μM]
p=0.002
p=0.0009
CFU / % Untreated
ITD-NPM1-2
Healthy Cells
Primary Cell Sample
C
ITD-13
log2FC
ATAC-Seq
DMSO
CBFβi
Rep1
Rep2
Motifs
AP-1
NFkB
ETS
RUNX
RNA-Seq logFC
Gained n = 265
Ranked by fold-difference Maintained; n = 16482
Lost n = 285
-/+ 1kb
D
FLT3-ITD specific genes
FLT3-ITD specific genes in RUNX1 module
log2FC RNA(CPM) CBFβi-Untreated
E
Module association of genes up-regulated after CBFβi
In module
Not in module
shRNA screen hit
shRNA screen multiple hits
RUNX1/NFIX/AP-1 cluster
EGR
FOX
CEBP
NF1
RUNX
AP-1

**Figure S6 (Related to Figure 6): Perturbation of RUNX1 with CBFβi in FLT3-ITD primary cells**
A: Representative images from proximity ligation assay (PLA) showing a time course of dissociation of the CBFβ::RUNX1 complex in primary FLT3-ITD (ITD-14) AML cells after treatment with 10 µM CBFβi. The red signal shows interactions between CBFβ and RUNX1 counterstained with DAPI (blue). Images show these interactions at 4 different time points. The scatter plot shows the number of red foci counted per cell in triplicate at the four different time points. B: Colony forming ability of primary FLT3-ITD AML and healthy cells after treatment with CBFβi. Significant p-values are indicated on the graph and were calculated using Student's t-test. C: Density plot of ATAC-Seq analysis (red) of a second primary FLT3-ITD patient cells (ITD-13) with and without CBFβi ranked against each other according to fold-change with the indicated TF motifs (black) at the open chromatin sites and the logFC RNA expression of the genes associated to the peaks present plotted alongside.
D: Unsupervised clustering of the fold change of gene expression in 3 different patients (ITD-12, ITD-13, ITD-14) after 24 h treatment with 10 µM CBFβi or 0.1% DMSO control. Left panel: FLT3-ITD AML specific genes. Right panel: FLT3-ITD AML specific genes in RUNX1 module.
E: Genes up-regulated in the RNA-seq data in 2 or more of the CBFbi treated patients. Heatmap shows the gene modules associated with each gene (black = associated, white = not). Genes which were single hits in the screen in 1 or more samples are in bold, if there were multiple hits in 1 or more samples they are in italics. Genes not in the RUNX1, AP1, CEBP, EGR1, FOX, NF1 module are not included in this data. Hierarchical clustering was performed to group genes in similar modules.

Figure S7

A

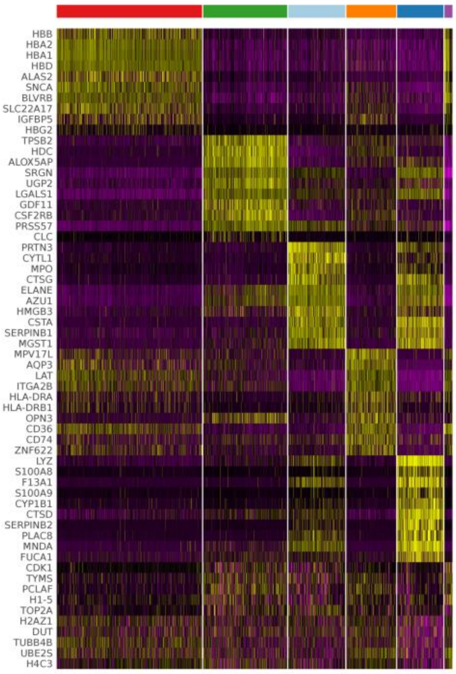


**No. of differentially expressed genes per cluster**

| Cluster ID | Up-Reg | Down-Reg |
|---|---|---|
| Early Progenitor | 139 | 271 |
| Eryth Prog - Quiescent | 80 | 121 |
| Eryth Prog - Cycling | 27 | 2 |
| Erythroid | 112 | 131 |
| Early Myeloid | 108 | 246 |
| Myeloid | 53 | 83 |

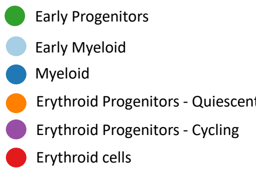


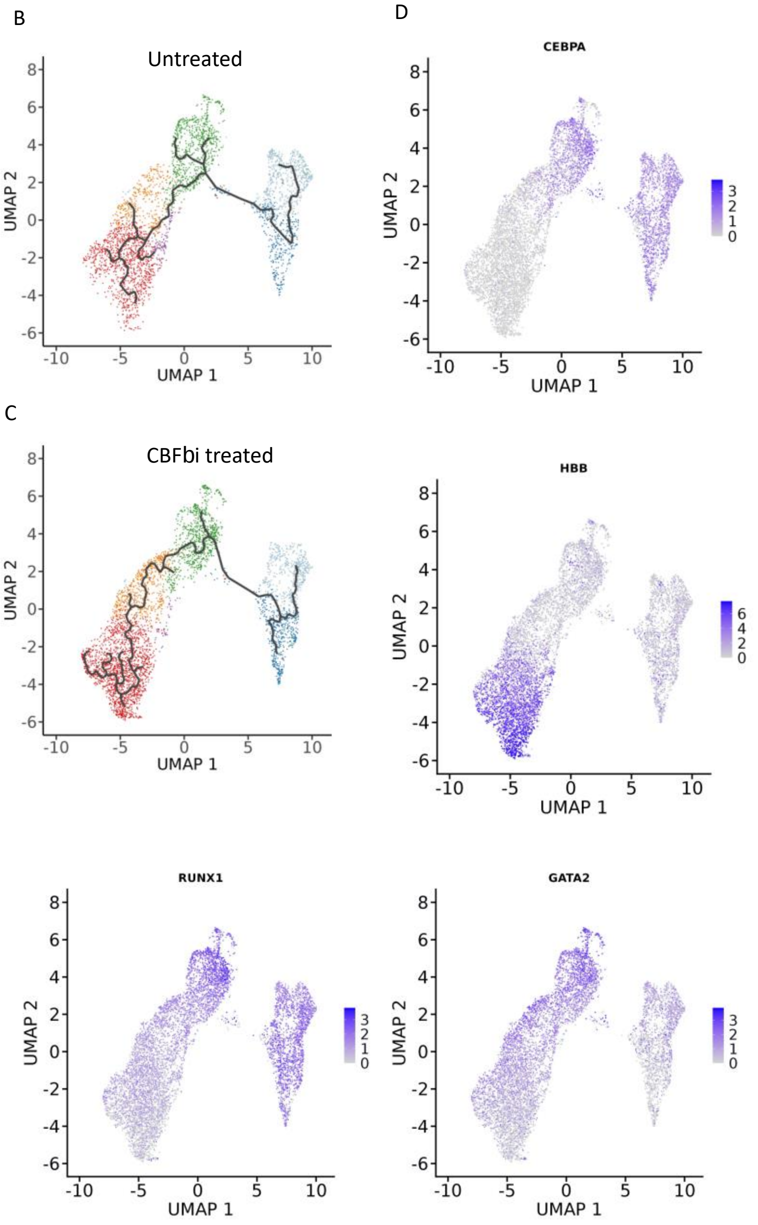


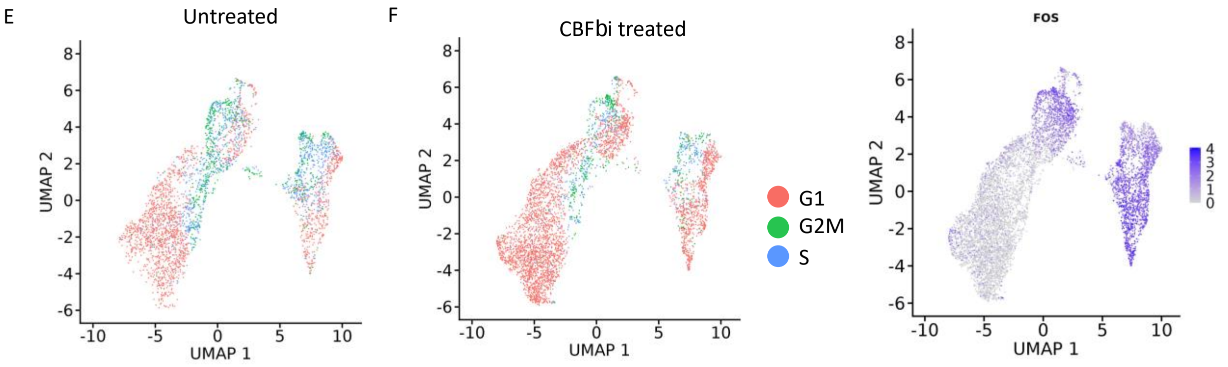

**Figure S7 (related to Figure 7): scRNA-Seq analysis of CBFβi treated FLT3-ITD AML patient cells**

A: Clustering analysis of scRNA data of the top 10 differentiallly expressed genes in each cluster defining the different cell types in the primary FLT3-ITD AML (ITD-12) culture. B, C: Pseudotime analysis of scRNA data without (B) and with (C) 10 µM CBFβi treatment. D: Expression of the indicated genes in the different cell types plotted onto the UMAP plots. E,F: UMAP plot depicting cell type in the indicated cell cycle phase without (E) and with (F) 10 µM CBFβi treatment.

### Supplementary Materials and Methods

#### Primary sample and PBSC processing

Human tissue was obtained with the required ethical approval from the National Health Service (NHS) National Research Ethics Committee. AML and PBSC samples used in this study were either surplus diagnostic samples or fresh samples obtained with specific consent from the subjects. AML samples were obtained from (1) the Haematological Malignancy Diagnostic Service (St James's Hospital, Leeds, UK), (2) the Centre for Clinical Haematology, Queen Elizabeth Hospital Birmingham, Birmingham, UK, or (3) the West Midlands Regional Genetics Laboratory, Birmingham Women's NHS Foundation Trust, Birmingham, UK. Mononuclear cells were purified on the same day they were received, and in most cases were also directly further purified using either CD34 or CD117 (KIT) magnetic antibodies as described[2]. For some samples with >92% blast cells, column purification was not performed. Mobilized PBSCs were provided by NHS Blood & Transplant, Leeds, UK, and NHS Blood & Transplant, Birmingham, UK.

#### Cell lines

For this study, we used two cell lines containing a FLT3-ITD mutation[3], MV4-11 (DMSZ, AC102) and MOLM14 (DSMZ, ACC 777). The cells were cultured in RPMI 1640 supplemented with 1% L-glutamine and 20% heat-inactivated FBS. For culture maintenance cells were split to $0.5x10^6$ cells/ml every 3 days to not exceed 1-2 x $10^6$ cells/ml. For the in vitro screen after sorting the media was also supplemented with 1% Penicillin/Streptomycin. HEK293T cells (DSMZ, ACC305) were used to produce lentivirus. These cells are cultured in HEPES-modified DMEM medium supplemented with 10% FBS, 4mM L-glutamine and 1mM sodium pyruvate. Cells were split using trypsin every 3 days to not exceed a confluency of 70%. All cells were cultured and treated in an incubator at 37C with 5% $CO_2$.

#### FLT3-ITD AML shRNA screen

##### Vector

The vector used in this study (now named pL40C) is described in detail in[4]. As described, the vector contains ampicillin resistance, a doxycycline-induced cassette that expresses the shRNA together with the fluorochrome dTomato and a constitutively expressed cassette containing the fluorochrome Venus.

#### shRNA oligo design

The shRNA oligos were designed using the informatic tool (https://felixfadams.shinyapps.io/miRN/) described previously[4,5]. 161 genes were included as targets and 3 shRNA oligos were designed per gene as described in the txt. As a positive control, FLT3 was included together with 10 NTC shRNA as negative controls. The oligos were ordered from Sigma Aldrich. Each oligo was 67 bp and was received with pre-mixed forward and reverse oligos at a concentration of 100 µM with desalt purification.

**shRNA library cloning**

The library of shRNA was produced following the process described in[4]. Briefly, the oligos were phosphorylated and annealed. Afterwards, all the oligos were pooled together. The vector was opened using the restriction enzyme BsmBI (Thermo, ER0451) following manufacturer recommendations. The opened plasmid was separated by running the product of the digestion in an agarose gel and extracting the DNA from the band using the Qiaquick Gel Extraction Kit (Qiagen, 28706) following manufacturer instructions. Plasmid and oligos were ligated using T4 DNA ligase kit (Thermo fisher, EL0011) with a molar ratio of 1:3 of vector and oligo. The ligation product was then transformed and amplified using XL-gold bacteria (Agilent, 200315) according to the manufacturer's protocol. After the transformation a maxiprep was perform to obtain high amounts of the cloned library, we used the EndoFree Plasmid Maxi Kit (Qiagen, 12362) and followed manufacturer's instructions.

**Lentivirus production**

The production of the lentivirus was done following the protocol described in Martinez et al[6].In summary, HEK293T cells were cultured in 15cm petri dishes up to a confluence of 50-60%. The library vectors and vectors for packaging and envelope (pMD2.G and pCMVΔR8.91) were mixed with special water (deionized water with 2.5 mM HEPES) and $CaCl_2$ 0.5 M solution. Next, the mix of the plasmid/special water and $CaCl_2$ was combined by dropwise addition o f HeBS. After incubation the solution was poured dropwise into the plates for transfection of the HEK293T cells. After one day the media is change, following 48h of incubation the supernatant containing the virus is collected, spin and freeze.

**Cell transduction**

MV4-11 and MOLM-14 cells were transduced following the protocol described in Martinez et al[6]. As a summary, cells at $10^6$ cells/ml were transduced with the pooled library shRNA lentivirus particles present in the supernatant using Polybrene at a final concentration of 8 µg/ml. Afterwards, the plate was centrifuged at 34C for 50 min at 900xg. After centrifugation, cells were incubated for 3 days. The transduction was performed at a low MOI (0.3 TU/cell) to produce a population of cells with one integration event per cell. Following lentiviral transduction, cells successfully carrying the fluorescent Venus constitutive expressed were purified using fluorescence activated cell sorting (FACS) in an ARIA II. Most cells that remained after selection carried a single copy of the inducible shRNA. Cells were then used to perform the screening as described.

***In vitro* screen**

For the *in vitro* screen we aimed for a coverage of 1000x of the library. Two different conditions were tested no doxycycline and doxycycline treatment. For transduced MOLM-14 the concentration of doxycycline (Sigma Aldrich, D5207) used was 500 ng/mL and 1µg/ml for transduced MV4-11. 5 million cells per condition were cultured at a concentration of $0.5x10^6$

cells/ml, split every 3 days to maintain that concentration, change the media and refresh the doxycycline. Cells were cultured for 15 passages and samples were collected at different time points. For obtaining DNA, cells were collected, spin and freeze in a pellet. The DNA was then isolated using the DNeasy Blood & Tissue Kit (Qiagen, 69504) following manufacturer's instructions.

**Mouse studies and PDX generation**

All mouse studies were carried out in accordance with UK Animals (Scientific Procedures) Act, 1986 under project licence P74687DB5 following approval from Newcastle University animal ethical review body (AWERB). Mice were housed in specific pathogen free conditions in individually ventilated cages with sterile bedding, water and diet (Irradiated RM3 breeding diet, SDS Ltd). All procedures were performed aseptically in a laminar flow hood.

NSG mice (NOD.Cg-Prkdcscid Il2rg tm1Wjl/SzJ) aged between 12 and 16 weeks, both sexes, from an in-house colony were used for PDX generation. They were transplanted intra-femorally with 1x10e6 patient or PDX cells under isoflurane anaesthetic and administered with subcutaneous NSAID analgesia (5 mg/kg subcutaneous Carprofen).

Mice were checked daily, weighed and examined at least once weekly to ensure good health. Endpoints for humane killing were pale extremities, hunched posture, 20% weight loss compared to highest previous weight or 10% weight loss for 3 consecutive days.

PDX cells were harvested from spleen and isolated by passing through a 50µM cell sieve (Falcon Corning). Cells were washed in PBS and stored frozen in 10%DMSO/90%FBS.

Mice used in the screen were Rag2$^{-/-}$Il2rg $^{-/-}$ 1293Balb/c (RG) mice (female) aged 8-10 weeks at study commencement.

**In vivo shRNA screen**

10 female mice Rag2$^{-/-}$Il2rg $^{-/-}$ 1293Balb/c (RG) from an in-house colony and aged 8-10 weeks were injected intra-venously with 50.000 MOLM-14 cells containing the shRNA library per mouse in a volume of 100µl. Mice were randomly assigned to 2 groups .One group was fed the normal RM3 diet and one doxycycline containing diet ( 823747 - CRM (E) + 625ppm Doxycycline (P) 1kg 25kG, SDS Ltd) ad libitum on the day of cell injection. Diet was replaced every 3 days and mouse health assessed daily. Mice were humanely killed 19-22 days after cell injection when a weak tail or hind legs were first detected. Cells were isolated from spleen as above. Cells were isolated from the bone marrow by crushing the leg and hip bones in PBS in a pestle and mortar, vortexing and passing the supernatant through a cell sieve. Engrafted cells were sorted by FACS (Aria II, BD) using Venus and dTomato fluorophore DNA was isolated using the DNeasy Blood & Tissue Kit (Qiagen, 69504) following manufacturer's instructions.

**Library preparation for shRNA screens**

PCR of genomic DNA was performed using ExTaq (Takara) using custom designed primers with Nextera i5 and i7 index sequences (supplementary table X) to amplify the mir30 insert containing the shRNA. Amplicons were electrophoresed on an agarose gel and DNA was purified using the QIAquick gel extraction kit (QIAGEN) according to manufacturer's instructions and further purified by ampure (Beckman Coulter). Samples were pooled and analysed on a Next Seq 2000 75 using a NextSeq 500/550 High output kit.

**FLT3-ITD cell line validations**

MV4-11 cells were transduced with lentiviral vectors and cultured as described above. For colony formation assays, MV4-11 cells were treated with 1 µg/ml doxycycline for 24 h prior to seeding at 5000 cells/ml in Methocult H4100 (StemCell Technologies) supplemented with Iscove's MDM (Merck) and FCS (Gibco) at a 2:2:1 ratio. 1 µg/ml doxycyline was added to the media and colonies were counted after 8 days.

For Western blot analysis of protein expression, MV4-11 cells were cultured for 3 days with 1 µg/ml of doxycycline added every 48 h. 12 µg of protein extracts in Laemmli buffer were run on a 4–20% gradient pre-cast gel (Bio-Rad) and transferred to nitrocellulose using Turbo transfer packs (Bio-Rad). Membranes were blocked with 10 % milk in TBS-T (10 mM Tris-HCl pH 7.5, 75 mM NaCl, 0.1% Tween-20) before being incubated at 4 °C overnight in 5 % milk TBS-T with primary antibody (αEGR1: 1:1000 (sc110 – SantaCruz), αNFIL3: 1:1000 (A302-606A, Bethyl), αRUNX1 (1:1000, 8758 cell signalling)). After washing in TBS-T, membranes were incubated in 5% milk TBS-T with HRP-conjugated anti-rabbit or anti-goat antibody (Cell Signalling Technologies) for 1 h at room temperature. After a further 3 washes in TBS-T, enhanced chemiluminescent reagent (Amersham) was applied and the blot was visualised using a GelDoc system (Bio-Rad). For loading controls, the membranes were stripped using Restore Stripping Buffer (Thermo Fisher Scientific) and GAPDH (ab8245; Abcam) was applied and visualised as above.

**Primary FLT3-ITD AML transcription factor perturbation experiments and colony forming assays**

Primary human AML blast cells were cultured on primary human mesenchymal stem cells (hMSCs). Primary hMSCs from "normal" bone marrow were cultured in alpha-MEM (Lonza) supplemented with 10 % fetal calf serum (Gibco) and 2 mM L-Glutamine (Gibco). 24 h prior to addition of primary AML cells hMSCs were seeded at 5000 cells/$cm^2$ in tissue culture plates pre-treated for 20 minutes with 0.1% glycine.

Primary human AML blasts were defrosted and cultured at 0.3-0.5 x $10^6$ cells/ml on hMSC feeders in alpha-MEM (Lonza) supplemented with 12.5 % fetal calf serum, 12.5 % horse serum, 100 U/ml penicillin/streptomycin, 2 mM L-Glutamine (all Gibco), 1 µM hydrocortisone (Merck) and 57.2 µM β-mercaptoethanol (Merck), 20 ng/ml IL-3, G-CSF and TPO (Pepro Tech) as described previously[7]. Cells were passaged to new feeders every 7 days. All cells were cultured and treated in an incubator at 37C with 5% $CO_2$.

**Inhibitor experiments in primary AML cells and healthy cells**

The DUSP 1/6 inhibitor BCI (Selleckchem) and FLT3-ITD inhibitor Quizartinib (Selleckchem) were dissolved to a 10 mM stock concentration in DMSO (Merck) on arrival. CBFβi (AI-14-91) and its control compound (AI-4-88)[8] were both dissolved to a 40 mM concentration in DMSO. Prior to dosing primary cells were cultured as described above for 7 days after defrost. Samples were then transferred to a 96 well plate previously prepared with hMSC feeders and the desired concentration of inhibitor was added to the media ("untreated" control was treated with 0.1% DMSO). Cells were then incubated with the inhibitors for 6 days before viability was assessed by counting cells on a haemocytometer after a 1:1 dilution with Trypan Blue (Merck) to differentiate alive and dead cells. For dose response curves IC50 was calculated using Graphpad prism software by performing non-linear regression (log[inhibitor] vs normalized response).

For colony formation assays – cells were treated with the inhibitor for 24h prior to seeding at a density of 5000 cells/ml in Methocult Express (StemCell Technologies). The inhibitor was also added to the colony medium at the same concentration. Colonies were counted after 12 days.

For NGS experiments – primary cells were treated with the desired concentration of inhibitor for 24 h prior to harvest with 0.1% DMSO as a control.

**Lenitviral transduction of primary AML cells and healthy cells**

pL40c shRNA were generated by cloning shRNAs (supplementary table X) into the pL40c vector. The dnFOS and dnCEBP inserts, originally generated by Charles Vinson (National Cancer Institute, Bethesda, MD, USA), were cloned into a pENTR backbone and then Gateway cloning was used to insert that into the Tet-on plasmid pCW57.1 (David Root, Addgene plasmid 41393).

For virus production, Human embryonic kidney 293T (HEK293T) cells were cultured in DMEM supplemented with 10% FCS, 2 mM L-glutamine, 100 U/ml penicillin, 100 mg/ml streptomycin and 0.11 mg ml–1 sodium pyruvate and were seeded to achieve 70–80% confluence at time of transfection. HEK293T cells were transfected using calcium phosphate co-precipitation of the five plasmids (LEGO-iG with TAT, REV, GAG/POL and VSV-G) at a mass ratio of 24 µg:1.2 µg:1.2 µg:1.2 µg:2.4 µg per 150 mm–diameter plate of cells. Viral supernatant was harvested after 24 h and subsequently every 12 h for 36 h before concentration with Centricon Plus-70 100-kDa filter (Millipore), using the manufacturer's instructions. Concentrated viral particles were stored at -70 °C before lentiviral transduction. Cell lines were transduced with concentrated virus in the presence of 8 µg/ml polybrene and 1x CD34 supplement (StemCell Technologies) by spinoculation at 1,500g for 50 min. After 12–16 h incubation at 37 °C, viral medium was exchanged for fresh medium. Cells were cultured for 3 days prior to treatment with 1.5 µg/ml doxycycline (Merck), with a further treatment after an additional 48 hours. After 3 days doxycycline treatment FACS was performed to isolate GFP+ (pCW57.1 dnFOS & dnCEBP) or Venus+/Tomato+ cells (pL40c-shRNA). For colony formation assays – sorted cells were seeded at 5000 cells/ml in Methocult Express (StemCell Technologies) with 1.5 µg/ml doxycycline and counted after 12 days.

**Mini-shRNA screen in healthy cells**

For shRNA mini screen in healthy cells, cells were not treated with doxycycline prior to FACS and Venus+ cells were collected. Cells were cultured for 12 days with 1.5 µl doxycycline added every 3 days. After 12 days DNA was extracted from cultured cells in the presence or absence of doxycycline and genomic DNA was extracted using the DNeasy blood and tissue kit (QIAGEN) according to manufacturer's instructions.

**ATAC-seq analysis of primary cells**

Omni ATAC-seq was performed as in Corces et al.[9]. Briefly, cells were washed in ATAC resuspension buffer (RSB) (10mM Tris-HCl pH7.5, 10mM NaCl and 3mM MgCl2) and then lysed for 3 minutes on ice in RSB buffer with 0.1% NP-40, 0.1% Tween-20. Then the cells were washed with 1ml of ATAC wash buffer consisting of RSB with 0.1% Tween-20. Then the nuclear pellet was resuspended in ATAC transposition buffer consisting of 25µl TD buffer and a concentration of Tn5 transposase enzyme (Illumina) related to the number of input cells, 16.5 µl PBS, 5 µl water, 0.1% tween-20 and 0.01% digitonin and then incubated on a thermomixer at 37°C for 30 minutes. The transposed DNA was then amplified by PCR amplification up to ½ of maximum amplification, as assessed by a qPCR side reaction. The library was purified using a QIAquick PCR cleanup kit (QIAGEN) followed by ampure (Beckman Coulter) and analysed on a Next Seq 2000 75 using a NextSeq 500/550 High output kit.

**RNA-seq of primary cells**

RNA was extracted from primary cells using a RNeasy Micro Plus kit (QIAGEN) where less than 50,000 cells were harvested, and a RNeasy Micro Plus kit (QIAGEN) for larger cell numbers. After quantification by nanodrop and QC using an Agilent RNA 6000 Pico Kit (Agilent, bioanalyser), libraries for next generation sequencing were prepared using the NEBnext Ultra II Directional RNA Library Prep Kit for Illumina (NEB) with the NEBNext® rRNA Depletion Kit v2 for low RNA input (<100 ng RNA), or the Total RNA Ribo-zero library preparation kit (with ribosomal RNA depletion) (Illumina) for higher RNA input. Libraries were quantified using the High Sensitivity DNA kit (Agilent) and Kapa Library Quantification kit (Roche) prior to paired end sequencing on a Next Seq 2000 (PE 75) with a NextSeq High 150 v2.5 kit.

**Proximity Ligation Assay (PLA ) of CBFb:RUNX1 interaction**

$1.5 \times 10^5$ cells were adhered to microscope slides using a Cytospin cytocentrifuge (Thermo Fisher Scientific) for 3 min at 800g and fixed in 4% formaldehyde (Pierce) for 15 min. Cells were permeabilised in 0.1% Triton X-100 and nonspecific staining was prevented by incubation in 3% bovine serum albumin. Anti-CBFβ (sc-56751; Santa Cruz Biotechnology at 1:100) and anti-RUNX1 (ab23980, Abcam) at 1:100 primary antibodies were applied for 1 hr at room temperature in PLA antibody diluent solution. Probes, ligation, and amplification solutions (Duolink; Sigma-Aldrich) were then applied at 37°C according to the manufacturer's instructions, and the slides were mounted in Duolink mounting medium with DAPI (Sigma-Aldrich). Slides were visualised using a Zeiss LSM 780 equipped with a Quasar spectral (GaAsP) detection system, using a Plan Achromat 40× 1.2 NA water immersion objective, Lasos 30 mW Diode 405 nm, Lasos 25 mW LGN30001 Argon 488, and Lasos 2 mW HeNe 594 nm laser lines. Images were acquired using Zen black version 2.1. Post-acquisition brightness and contrast adjustment was performed uniformly across the entire image.

**RUNX1 ChIP seq from ITD-14 patient cells**

Primary AML cells were cultured on hMSC feeders as described above in the following media: SFEMII (StemCell Technologies), 1 µM UM729 (StemCell Technologies), 750 nM StemReginin 1 (StemCell Technologies) supplemented with 150 ng/ml SCF, 100 ng/ml TPO, 10 ng/ml IL-3, 10 ng/ml G-CSF (Perpro tech). After 1 passage (1 week) in culture cells were treated for 24 h with 10 µM CBFβi or 0.1 % DMSO in the absence of UM729 and StemReginin 1. 2 million cells were harvested and crosslinked in 1% formaldehyde solution (methanol-free from Pierce, Thermo Scientific). 400 mM of glycine (Merck) was added, and cells were washed twice with PBS (Merck) after which pellets were frozen at -80 °C.

Crosslinked cells were resuspended at $1x10^7$ cells/ml in Buffer A (10 mM HEPES, 10 mM EDTA, 0.5 mM MEGTA, 0.25 % Triton- 100, 1x complete mini protease inhibitor cocktail (PIC) (Merck) pH 8.0) and incubated at 4 °C for 10 minutes prior to centrifugation at 500 xG for 10 min. This step was repeated with Buffer B (10 mM HEPES, 200 mM NaCl, 1 mM EDTA, 0.5 mM MEGTA, 0.01 % Triton-X 100, 1x PIC, pH 8.0) and after centrifugation 2 x $10^6$ cells were resuspended in 300 µl IP buffer I (25 mM Tris-HCl, 150 mM NaCl, 2 mM EDTA, 1 % Triton-X 100, 0.25 % SDS, 1x PIC, pH 8.0) and sonicated using a Diagenode Bioruptor Pico sonicator for 4 cycles (30 sec on 30 sec off) before centrifugation for 10 min at 16,000 xG. The supernatant was then collected and 600 µl IP Buffer II (25 mM Tris-HCl, 150 mM NaCl, 2 mM EDTA, 1 % Triton-X 100, 7.5 % Glycerol, 1x PIC, pH 8.0) was added prior to immunoprecipitation.

For immunoprecipitations 15 µl of Dynabeads-Protein G were washed twice with 500 µl 50 mM citrate phosphate buffer pH 5 and resupended in 15 µl citrate phosphate buffer with 4 µg anti-RUNX1 antibody (ab23980, Abcam) and 0.5% acetyl-BSA before incubation at 4 °C for 2 h. After incubation, dynabeads were washed with 500 µl pH 5 citrate phosphate buffer and resuspended in 15 µl citrate phosphate buffer with 0.5 % BSA before 555 µl of sonicated chromatin was added and incubated at 4 °C for ~ 16 h.

After the incubation the dynabeads are washed sequentially with 500 µl of: Wash buffer 1 (20 mM Tris-HCl, 150 mM NaCl, 2 mM EDTA, 1% Triton-X 100, 0.1% SDS, pH 8.0) once, Wash buffer 2 (20 mM Tris-HCl, 500 mM NaCl, 20 mM EDTA, 1 % Triton-X 100, 0.1 % SDS, pH 8) twice, LiCl buffer (10 mM Tris-HCl,250 mM LiCl, 1 mM EDTA, 0.5 % NP-40, 0.5 % Na-deoxycholate, pH 8.0) once, TE/NaCl wash buffer (10 mM Tris-HCl, 50 mM NaCl, 1 mM EDTA, pH 8.0) twice. After these washes DNA was eluted from the dynabeads using 100 µl elution buffer (100 mM $NaHCO_3$, 1% SDS). 200 mM NaCl and 500 µg/ml proteinase K were added to the eluant and the sample was reverse crosslinked at 65 °C for >4h. DNA was then purified by ampure (1.8x). Libraries for next generation sequencing were prepared using a Kapa Hyper prep kit (Roche) according to manufacturers instructions. 16 cycles of PCR amplification were used and 200-450 bp fragments were size selected by gel electrophoresis. Libraries were validated by qPCR and quantified using the High Sensitivity DNA kit (Agilent) and Kapa Library Quantification kit (Roche) prior to sequencing on a Nextseq 2000 75 using a NextSeq 500/550 High output kit.

**Single cell treatment scRNA-Seq analysis of CBFβi treated FLT3-ITD+ AML**

Primary AML cells for scRNA-seq were cultured on hMSC feeders as described above in the following media: SFEMII (StemCell Technologies), 1 µM UM729 (StemCell Technologies), 750 nM StemReginin 1 (StemCell Technologies) supplemented with 150 ng/ml SCF, 100 ng/ml TPO, 10 ng/ml IL-3, 10 ng/ml G-CSF (Perpro tech). After 1 passage (1 week) in culture cells were treated for 24 h with 10 µM CBFβi or 0.1 % DMSO in the absence of UM729 and StemReginin 1. After treatment cells were sorted for CD45 using magnetic beads (Miltenyi Biotec). Cells were loaded on a Chromium Single Cell Instrument (10X Genomics), to recover 5000 single cells. Library generation was performed using the Chromium single cell 3' library and gel bead kit v3.1. Illumina sequencing was performed on a NovaSeq 6000 S1 run in paired-end mode for 150 cycles at a depth of 20000 reads per cell.

**Bulk RNA-Seq data analysis**

Raw paired-end reads were trimmed to remove low-quality sequences and adaptors using Trimmomatic v0.39[10]. Reads were then aligned to the human genome (version hg38) using HISAT2 v2.2.1[11] with default settings. Counts were generated with featureCounts v2.0.1[12] using gene models from ensembl as the reference transcriptome. Differential gene expression analysis was carried out using Limma-Voom v3.50.3[13] in R v4.1.2.

**Single-Cell RNA-Seq analysis**

Fastq files from single-cell sequencing experiments were aligned to the human genome (version hg38) using the count function in CellRanger v5.0.1 from 10x genomics[14] using gene models from ensembl as the reference transcriptome. Analysis was then carried out using the Seurat package v4.3.0[15] in R v4.1.2. Cells from CBFbi treated and untreated samples were filtered to remove cells with less than 500 and more than 6000 detected genes, as well as cells with more than 20% of reads aligned to mitochondrial transcripts. The filtered cells were then combined into a single dataset for downstream analysis. UMI counts were normalized using the NormalizeData function with default settings. The cell cycle stage for each cell was inferred using the CellCycleScoring function in Seurat. This score was then used to remove the possible effect of cell cycle stage on the analysis by linear regression using the ScaleData function. Clustering was then carried out using the FindNeighbors and FindClusters commands, using the top 20 principal components and a cluster resolution value of 0.25. Differential gene expression analysis was carried out for each single cell cluster, comparing CBFBi treated cells to untreated cells using the FindMarkers command. A gene with a log2 fold-change of at least 0.25 and an adjusted p-value less than 0.1 were considered to be differentially expressed. Kyoto Encyclopaedia of Genes and Genomes (KEGG) pathway analysis was then carried out on the sets of differentially expressed genes using the ClueGO package v2.5.0[16] in Cytoscape v3.9.1[17]. Cell trajectory (pseudotime) analysis was carried out using Monocle3 v1.3.1[18]. To do this, the processed data from Seurat was first divided into two objects corresponding to CBFBi treated and untreated samples. These were then imported into Monocle using the as.cell_data_set function in SeuratWrappers. Single cell trajectories were calculated using the cluster_cells and learn_graph commands in Monocle. Pseudotime was then calculated by rooting the trajectory at the earliest point of the inferred trajectory that occurred in the early progenitor cells.

**shRNA data analysis**

To calculate read counts from shRNA experiments, 75bp single-end reads in fastq format were first processed to remove the first and last 25bp from each sequence, corresponding to the regions flanking the shRNA sequence that are common across all reads. The shRNA sequences were then compared to the library of oligonucleotide sequences used in the experiment, allowing for only a single base mismatch. Read counts were normalized using upper-quartile normalization using the edgeR package v3.36.0[19] in R v4.1.2. To calculate fold-changes between doxycycline induced and non-induced cells, the normalized counts were fitted to a generalized linear model using edgeR. A shRNA sequence was deemed to have been lost if it had a log2 fold-change less then -1 between induced and non-induced samples.

**ATAC-Seq data analysis**

Single-end reads from ATAC-Seq experiments were processed to remove low-quality sequences and Nextera ATAC adaptors using Trimmomatic. Reads were then aligned to the human genome (version hg38) using Bowtie2 v2.2.5[20] with the option --very-sensitive-local. Potential PCR duplicates were identified and removed from alignments using Picard MarkDuplicates v2.26.10 (http://broadinstitute.github.io/picard). Peaks were called using MACS2 v2.2.7.1[21] with the parameters --nomodel -B --trackline. The resulting peaks were then filtered to remove any peak with a peak height less than 10 or were found in the hg38 blacklist[22] Where replicates were available, only peaks that passed these filters in both replicates were retained. A peak union was then created for each set of experiments by first extending the peak region by 200bp either side of the peak summit. Overlapping peaks were then combined using the merge function in bedtools v2.30.0[23]. The distance between the peak summit and the closest gene was then calculated using the annotatePeaks.pl function in Homer v4.9.1[24]. A peak was classified as distal if it was at least 1.5kb from the nearest transcriptional start site (TSS), and as promoter-proximal otherwise. Distal and promoter-proximal peaks were treated separately in downstream analyses.

Differential peak analysis was carried out by first counting the number of reads aligned to each peak using featureCounts. These were then normalized as counts-per-million using the edgeR package in R. In cases where replicates were available, fold-differences and statistical values were calculated using Limma-Voom. Where only single experiments were available, a simple fold-difference was calculated by subtracting the log2-normalized count from the treatment sample from the control (NTC, empty-vector as appropriate). Read density plots were created by first ranking peaks according to fold-difference. The read counts were then retrieved in a 2kb window centered on the peak summit using the annotatePeaks.pl function in Homer with the options -size 2000 -hist 10 -ghist -bedGraph with the bedGraph files produced by MACS2 as input. These were then plotted as a heatmap using Java TreeView v1.1.6r4[25]. Motif enrichment analysis was carried out in the set of gained and lost peaks using the findMotifsGenome.pl function in Homer with the options -size 200 -noknown.

**ChIP-Seq data analysis**

Reads were trimmed to remove low-quality sequences using Trimmomatic and aligned to the human genome (version hg38) using Bowtie2 with the --very-sensitive-local paramter. Potential PCR duplicates were identified and removed from alignments using Picard MarkDuplicates. Peaks were then called using MACS2 with the settings -B --trackline. Read density plots were created by first ranking peaks by tag count, and then retrieving the read density in a 2kb region centered on the peak summit using the annotatePeaks.pl function in Homer. These were then shown as a heatmap in Java TreeView.

**Re-analysis of public DNaseI-Seq data**

DNaseI-Seq data from FLT3-ITD, FLT3-ITD + NPM1 and healthy PBSCs from Assi et al. (2019) were downloaded from GEO using the accession number GSE108316. Reads were then trimmed using Trimmomatic, and aligned to the human genome using Bowtie2 using the parameter --very-sensitive-local. Peaks were called using MACS2 with the options --keep-dup all --nomodel -q 0.0005 --call-summits -B --trackline. In order to ensure that peak coordinates were accurate and representative of all patients, alignments were merged to create a single dataset. Peak calling was then repeated on this dataset and the resulting peak coordinates were used as the reference peak positions for all further analysis. The peaks from each sample were filtered to remove peaks in the hg38 blacklist, and only peaks that were found in at least 50% of all patients from their respective groups (ITD, ITD-NPM1 or PBSC) were retained for further analysis. Differential peak analysis was carried out by first classifying peaks as either distal, or as promoter-proximal as described for the ATAC-Seq data above and processed separately. The average read count in a 400bp window centered on the peak summit was then retrieved using the annotatePeaks.pl function in Homer with the options -size 400 -bedGraph and using the bedGraph files produced by MACS2. These counts were then normalized to the average read count across samples. The average read count for each group (ITD/ITD-NPM1 or PBSC) was then calculated, and further log2-transformed as log2(average read count + 1). A peak was deemed to be specific to a group if it had a fold-difference greater than 3 in either the ITD or ITD-NPM1 groups compared to healthy PBSCs. DNaseI footprinting was carried out on the merged alignments from ITD, ITD-NPM1 and PBSCs using the wellington algorithm v0.2.0[26] using the options -fdrlimit -5 -fp 11,32,2. The resulting set of footprints were then combined using the bedtools merge command.

**Construction of Gene Regulatory Networks**

Gene Regulatory Networks (GRNs) were made using custom Python scripts that have been made publicly available (see code and data availability section). To construct ITD/ITD-NPM1 and PBSC specific networks, we first identified sets of DNaseI Hypersensitive Sites (DHSs) from Assi et al.[1] that had a fold-difference greater than 3 between cell types. The genomic positions of transcription factor binding motifs were then retrieved from within these sites using the annotatePeaks.pl function in Homer and exported as BED files using the -mbed option. To ensure that the motif sequences used in each of these networks were constant, we used the set of probability weight matrices that were defined in Assi et al.[1] and have been made available for further use (see code and data availability section). These motif positions were then further refined by only keeping those that were found within DNaseI footprints. To ensure that DHSs were assigned to the correct gene, we used processed promoter-capture HiC data from FLT3-ITD patients and healthy CD34 positive cells that were analysed by Assi et al.[1]. In cases where no HiC annotation was present for a given DHS, the peak was instead assigned to the closest gene. To ensure that only genes that were actually expressed in our data were included in the GRN, we used RNA-Seq data from Assi et al.[1] which includes data from the same set of patients that were used to construct the GRN. These were processed as described above, and only genes that were expressed with a Fragments Per Kilobase per Million mapped reads (FPKM) value greater than 1 in either ITD/ITD-NPM1 patients or healthy PBSCs were included. A network was then constructed for each set of specific and shared DHSs where transcription factor genes are represented by nodes, and the presence of a footprinted binding motif in a DHS targeting a gene is represented by a directed edge. Members of the same transcription factor families are known to bind to highly similar or identical motifs[27] making the definitive identification of specific transcription factor genes from motif data alone difficult. To account for this in our GRNs, we grouped members of the same family into groups with one binding motif representing the entire set of transcription factor genes. The most highly expressed member of that family was used as the source node. The GRN graph was then exported as a JSON file and visualised using Cytoscape.

**Transcription Factor module similarity**

In order to measure if the sets of genes that were targeted by different transcription factor families were similar, we first extracted the module for each TF family. A module here is defined as the complete set of target genes from the GRN and includes both transcription factor and non-transcription factor genes (figure 3A). The overlap of these modules was then measured using the Jaccard similarity index using the formula

$$Jaccard\ Index = \frac{|A \cap B|}{|A \cup B|}$$

Where A and B are the sets of genes for two different transcription factor modules. This was calculated for each pair of transcription factor modules, resulting in a matrix of Jaccard Index values. These were then hierarchically clustered using complete linkage of the Euclidean distance in R and shown as a heatmap (figure 3B-E).

**Supplementary References**


1. Assi, S.A., Imperato, M.R., Coleman, D.J.L., Pickin, A., Potluri, S., Ptasinska, A., Chin, P.S., Blair, H., Cauchy, P., James, S.R., et al. (2019). Subtype-specific regulatory network rewiring in acute myeloid leukemia. Nat Genet *51*, 151-162. 10.1038/s41588-018-0270-1.
2. Cauchy, P., James, S.R., Zacarias-Cabeza, J., Ptasinska, A., Imperato, M.R., Assi, S.A., Piper, J., Canestraro, M., Hoogenkamp, M., Raghavan, M., et al. (2015). Chronic FLT3-ITD Signaling in Acute Myeloid Leukemia Is Connected to a Specific Chromatin Signature. Cell Rep *12*, 821-836. 10.1016/j.celrep.2015.06.069.
3. Quentmeier, H., Reinhardt, J., Zaborski, M., and Drexler, H.G. (2003). FLT3 mutations in acute myeloid leukemia cell lines. Leukemia *17*, 120-124. 10.1038/sj.leu.2402740.
4. Adams, F.F., Heckl, D., Hoffmann, T., Talbot, S.R., Kloos, A., Thol, F., Heuser, M., Zuber, J., Schambach, A., and Schwarzer, A. (2017). An optimized lentiviral vector system for conditional RNAi and efficient cloning of microRNA embedded short hairpin RNA libraries. Biomaterials *139*, 102-115. 10.1016/j.biomaterials.2017.05.032.
5. Adams, F.F., Hoffmann, T., Zuber, J., Heckl, D., Schambach, A., and Schwarzer, A. (2018). Pooled Generation of Lentiviral Tetracycline-Regulated microRNA Embedded Short Hairpin RNA Libraries. Hum Gene Ther Methods *29*, 16-29. 10.1089/hgtb.2017.182.
6. Martinez-Soria, N., McKenzie, L., Draper, J., Ptasinska, A., Issa, H., Potluri, S., Blair, H.J., Pickin, A., Isa, A., Chin, P.S., et al. (2018). The Oncogenic Transcription Factor RUNX1/ETO Corrupts Cell Cycle Regulation to Drive Leukemic Transformation. Cancer Cell *34*, 626-642.e628. 10.1016/j.ccell.2018.08.015.
7. van Gosliga, D., Schepers, H., Rizo, A., van der Kolk, D., Vellenga, E., and Schuringa, J.J. (2007). Establishing long-term cultures with self-renewing acute myeloid leukemia stem/progenitor cells. Exp Hematol *35*, 1538-1549. 10.1016/j.exphem.2007.07.001.
8. Illendula, A., Gilmour, J., Grembecka, J., Tirumala, V.S.S., Boulton, A., Kuntimaddi, A., Schmidt, C., Wang, L., Pulikkan, J.A., Zong, H., et al. (2016). Small Molecule Inhibitor of CBFβ-RUNX Binding for RUNX Transcription Factor Driven Cancers. EBioMedicine *8*, 117-131. 10.1016/j.ebiom.2016.04.032.
9. Corces, M.R., Trevino, A.E., Hamilton, E.G., Greenside, P.G., Sinnott-Armstrong, N.A., Vesuna, S., Satpathy, A.T., Rubin, A.J., Montine, K.S., Wu, B., et al. (2017). An improved ATAC-seq protocol reduces background and enables interrogation of frozen tissues. Nat Methods *14*, 959-962. 10.1038/nmeth.4396.
10. Bolger, A.M., Lohse, M., and Usadel, B. (2014). Trimmomatic: a flexible trimmer for Illumina sequence data. Bioinformatics *30*, 2114-2120. 10.1093/bioinformatics/btu170.
11. Kim, D., Paggi, J.M., Park, C., Bennett, C., and Salzberg, S.L. (2019). Graph-based genome alignment and genotyping with HISAT2 and HISAT-genotype. Nature Biotechnology *37*, 907-915. 10.1038/s41587-019-0201-4.
12. Liao, Y., Smyth, G.K., and Shi, W. (2014). featureCounts: an efficient general purpose program for assigning sequence reads to genomic features. Bioinformatics *30*, 923-930. 10.1093/bioinformatics/btt656.

13. Law, C.W., Chen, Y., Shi, W., and Smyth, G.K. (2014). voom: Precision weights unlock linear model analysis tools for RNA-seq read counts. Genome Biol *15*, R29. 10.1186/gb-2014-15-2-r29.
14. Zheng, G.X., Terry, J.M., Belgrader, P., Ryvkin, P., Bent, Z.W., Wilson, R., Ziraldo, S.B., Wheeler, T.D., McDermott, G.P., Zhu, J., et al. (2017). Massively parallel digital transcriptional profiling of single cells. Nat Commun *8*, 14049. 10.1038/ncomms14049.
15. Hao, Y., Hao, S., Andersen-Nissen, E., Mauck, W.M., 3rd, Zheng, S., Butler, A., Lee, M.J., Wilk, A.J., Darby, C., Zager, M., et al. (2021). Integrated analysis of multimodal single-cell data. Cell *184*, 3573-3587.e3529. 10.1016/j.cell.2021.04.048.
16. Bindea, G., Mlecnik, B., Hackl, H., Charoentong, P., Tosolini, M., Kirilovsky, A., Fridman, W.H., Pagès, F., Trajanoski, Z., and Galon, J. (2009). ClueGO: a Cytoscape plug-in to decipher functionally grouped gene ontology and pathway annotation networks. Bioinformatics *25*, 1091-1093. 10.1093/bioinformatics/btp101.
17. Shannon, P., Markiel, A., Ozier, O., Baliga, N.S., Wang, J.T., Ramage, D., Amin, N., Schwikowski, B., and Ideker, T. (2003). Cytoscape: a software environment for integrated models of biomolecular interaction networks. Genome Res *13*, 2498-2504. 10.1101/gr.1239303.
18. Qiu, X., Mao, Q., Tang, Y., Wang, L., Chawla, R., Pliner, H.A., and Trapnell, C. (2017). Reversed graph embedding resolves complex single-cell trajectories. Nat Methods *14*, 979-982. 10.1038/nmeth.4402.
19. Robinson, M.D., McCarthy, D.J., and Smyth, G.K. (2010). edgeR: a Bioconductor package for differential expression analysis of digital gene expression data. Bioinformatics *26*, 139-140. 10.1093/bioinformatics/btp616.
20. Langmead, B., and Salzberg, S.L. (2012). Fast gapped-read alignment with Bowtie 2. Nat Methods *9*, 357-359. 10.1038/nmeth.1923.
21. Zhang, Y., Liu, T., Meyer, C.A., Eeckhoute, J., Johnson, D.S., Bernstein, B.E., Nusbaum, C., Myers, R.M., Brown, M., Li, W., and Liu, X.S. (2008). Model-based analysis of ChIP-Seq (MACS). Genome Biol *9*, R137. 10.1186/gb-2008-9-9-r137.
22. Amemiya, H.M., Kundaje, A., and Boyle, A.P. (2019). The ENCODE Blacklist: Identification of Problematic Regions of the Genome. Sci Rep *9*, 9354. 10.1038/s41598-019-45839-z.
23. Quinlan, A.R., and Hall, I.M. (2010). BEDTools: a flexible suite of utilities for comparing genomic features. Bioinformatics *26*, 841-842. 10.1093/bioinformatics/btq033.
24. Heinz, S., Benner, C., Spann, N., Bertolino, E., Lin, Y.C., Laslo, P., Cheng, J.X., Murre, C., Singh, H., and Glass, C.K. (2010). Simple combinations of lineage-determining transcription factors prime cis-regulatory elements required for macrophage and B cell identities. Mol Cell *38*, 576-589. 10.1016/j.molcel.2010.05.004.
25. Saldanha, A.J. (2004). Java Treeview--extensible visualization of microarray data. Bioinformatics *20*, 3246-3248. 10.1093/bioinformatics/bth349.
26. Piper, J., Elze, M.C., Cauchy, P., Cockerill, P.N., Bonifer, C., and Ott, S. (2013). Wellington: a novel method for the accurate identification of digital genomic footprints from DNase-seq data. Nucleic Acids Res *41*, e201. 10.1093/nar/gkt850.
27. Assi, S.A., Bonifer, C., and Cockerill, P.N. (2019). Rewiring of the Transcription Factor Network in Acute Myeloid Leukemia. Cancer Inform *18*, 1176935119859863. 10.1177/1176935119859863.
